# CIP2A is a prime synthetic-lethal target for BRCA-mutated cancers

**DOI:** 10.1101/2021.02.08.430060

**Authors:** Salomé Adam, Silvia Emma Rossi, Nathalie Moatti, Mara De Marco Zompit, Timothy F. Ng, Alejandro Álvarez-Quilón, Jessica Desjardins, Vivek Bhaskaran, Giovanni Martino, Dheva Setiaputra, Sylvie M. Noordermeer, Toshiro K. Ohsumi, Nicole Hustedt, Rachel K. Szilard, Natasha Chaudhary, Meagan Munro, Artur Veloso, Henrique Melo, Shou Yun Yin, Robert Papp, Jordan T.F. Young, Michael Zinda, Manuel Stucki, Daniel Durocher

**Affiliations:** Lunenfeld-Tanenbaum Research Institute, Mount Sinai Hospital, 600 University Avenue, Toronto, ON, M5G 1X5, Canada; Department of Gynecology, University Hospital and University of Zurich, Wagistrasse 14, 8952 Schlieren, Switzerland; Department of Molecular Genetics, University of Toronto, 1 King’s College Circle, Toronto, ON, M5S 1A8, Canada; Repare Therapeutics, 7210 Frederick-Banting, Suite 100, St-Laurent, QC, H4S 2A1, Canada

**Author notes:** These authors contributed equally to this work (Salomé Adam, Silvia Emma Rossi). Current address: Department of Human Genetics, Leiden University Medical Center, Leiden, The Netherlands (Sylvie M. Noordermeer). Current address: Ridgeline Therapeutics, Hochbergerstrasse 60C, CH-4057 Basel, Switzerland.

## Abstract

*BRCA1/2*-mutated cancer cells must adapt to the genome instability caused by their deficiency in homologous recombination. Identifying and targeting these adaptive mechanisms may provide new therapeutic strategies. Here we present the results of genome-scale CRISPR/Cas9-based synthetic lethality screens in isogenic pairs of BRCA1- and BRCA2-deficient cells that identified the gene encoding CIP2A as essential in a wide range of *BRCA1*- and *BRCA2*-mutated cells. Unlike PARP inhibition, CIP2A-deficiency does not cause accumulation of replication-associated DNA lesions that require homologous recombination for their repair. CIP2A is cytoplasmic in interphase but, in mitosis, accumulates at DNA lesions as part of a complex with TOPBP1, a multifunctional genome stability factor. In BRCA-deficient cells, the CIP2A-TOPBP1 complex prevents lethal mis-segregation of acentric chromosomes that arises from impaired DNA synthesis. Finally, physical disruption of the CIP2A-TOPBP1 complex is highly deleterious in BRCA-deficient cells and tumors, indicating that targeting this mitotic chromosome stability process represents an attractive synthetic-lethal therapeutic strategy for *BRCA1*- and *BRCA2*-mutated cancers.

---

The BRCA1 and BRCA2 proteins promote the repair of replication-associated DNA damage by homologous recombination (HR) ^1^. Acute inactivation of BRCA2 impedes completion of DNA replication ^2^, which is associated with rampant chromosome segregation defects and cell lethality. This phenotype is likely to be shared by BRCA1 since its loss also causes cellular lethality ^3^. These observations suggest that during their evolution towards the malignant phenotype, cells with inactivating mutations in *BRCA1* and *BRCA2* adapt to the replication-associated problems caused by HR deficiency. Identifying the mechanisms that endow cells to complete chromosome duplication and segregation without active HR may thus provide new opportunities for therapeutic intervention. The targeting of these adaptive mechanisms should display efficacy and toxicity profiles that are distinct from genotoxic chemotherapy or poly(ADP-ribose) polymerase (PARP) inhibition, since these latter therapies instead act by increasing the load of DNA lesions that engage HR-dependent DNA repair ^4^.

## The CIP2A-BRCA synthetic lethality

To identify a complement of genes that is essential for the viability of HR-deficient cells, we carried out genome-scale dropout CRISPR-based synthetic lethal screens in isogenic pairs of *BRCA1*- and *BRCA2*-mutated cells in the human RPE1-hTERT (immortalized retinal epithelium) and DLD1 (colon adenocarcinoma) backgrounds, respectively (Fig. S1A). We reasoned that genetic interactions common to the loss of BRCA1 and BRCA2 in two cell lines of different origins would have the potential to uncover universal vulnerabilities to the loss of HR.

The screens were carried out with the TKOv2 (BRCA1 screen) or TKOv3 (BRCA2 screen) single-guide (sg) RNA libraries and were analyzed with a custom-built analysis pipeline, called CRISPRCount Analysis (CCA), dedicated to the identification of synthetic-lethal genetic interactions (see methods), defined here as genes essential for the fitness of a mutated cell line (in this case *BRCA1*^−/−^ or *BRCA2*^−/−^) but not of their isogenic wild type counterparts. CCA identified 55 and 50 genes that selectively impaired fitness in the *BRCA1*- or *BRCA2*-mutated cells, respectively (Fig. 1A, Table S1 and Datasets S1-S2). The top 10 genes common to both screens were *APEX1*, *APEX2*, *CHD1L*, *CHTF8*, *CIP2A*, *DSCC1, DDIAS, PARP1, SLC25A28* and *XRCC1* (Fig. 1A). Of these, *PARP1* and *APEX2* are known to display robust synthetic lethal interactions with HR deficiency when depleted or inhibited ^5–8^. CHD1L was also recently shown to promote survival to PARP inhibition and to impair fitness of BRCA-deficient cells ^9,10^. Other genes with known synthetic lethal interactions with *BRCA1/2* such as *POLQ ^11,12^* or the RNase H2-coding genes ^6,9^ were hits in only one of the two cell lines (Table S1).

**Fig. 1.**
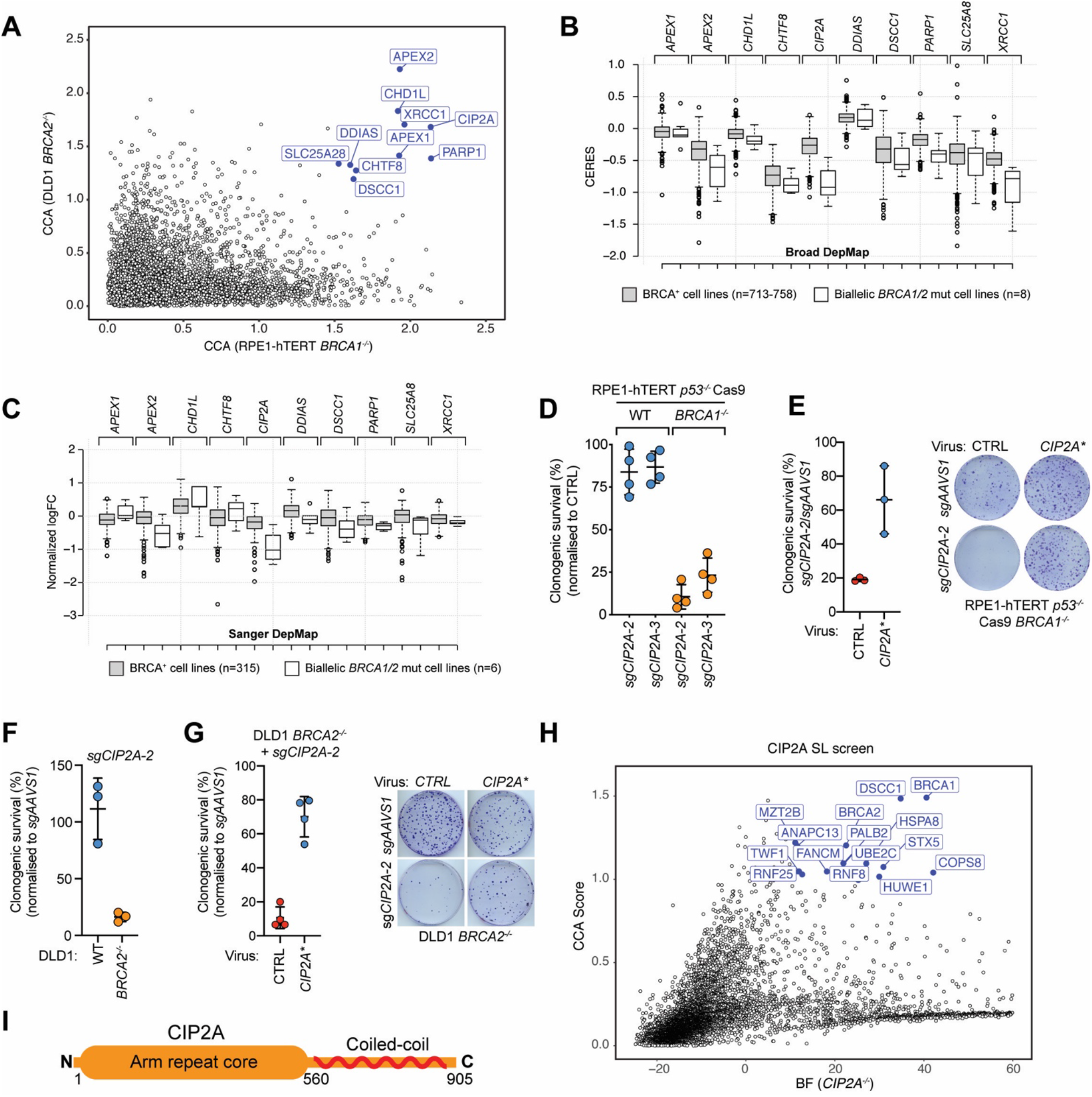
*CIP2A* loss is synthetic-lethal with *BRCA1-* or *BRCA2-*deficiency. (**A**) Scatter plot of CCA scores for the CRISPR synthetic lethality screens in *BRCA1^−/−^* and *BRCA2^−/−^* cells. Highlighted in blue are the top 10 genes common to both screens. (**B, C**) Boxplots of essentiality scores for the indicated genes derived from the Broad (B) and Sanger (C) DepMap projects. Cell lines were grouped according to whether or not they harbored biallelic inactivating mutations in *BRCA1* or *BRCA2.* Center lines show the medians; box limits indicate the 25th and 75th percentiles; whiskers extend 1.5 times the interquartile range, outliers are represented by dots. See Fig. S1B for statistical analysis of the results. (**D**) Clonogenic survival of RPE1-hTERT *p53*^−/−^ Cas9 wild-type (WT) and *BRCA1^−/−^* cells expressing the indicated *CIP2A*-targeting sgRNAs or transduced with control lentivirus (CTRL: either empty virus or virus with sgRNA targeting *AAVS1*). Data was normalized to the plating efficiency of the control virus. Data are shown as mean ± S.D. (n=4). (**E**) Reintroduction of a sgRNA-resistant *CIP2A* transgene (CIP2A*) rescues lethality of RPE1-hTERT *p53*^−/−^ Cas9 *BRCA1^−/−^* cells caused by *sgCIP2A-2*. Data are shown as mean ± S.D. (n=3). (**F**) Clonogenic survival of DLD1 wild-type (WT) and *BRCA2^−/−^* cells expressing *CIP2A*- or *AAVS1*-targeting sgRNAs. Data was normalized to the plating efficiency of cells expressing *sgAAVS1*. Data are shown as mean ± S.D. (n=4). (**G**) Reintroduction of a sgRNA-resistant *CIP2A* transgene rescues lethality in DLD1 *BRCA2^−/−^* cells caused by *sgCIP2A-2*. Data are shown as mean ± S.D. (n=3). (**H**) Scatter plot of CCA scores (y-axis) and Bayes Factor (BF) values derived from BAGEL2 (x-axis, for *CIP2A^−/−^* cell line) for the *CIP2A* isogenic synthetic lethal screen. (**I**) Schematic representation of the CIP2A protein.

To identify genetic interactions with highest relevance to the tumor setting, we analyzed the results of two large-scale studies of genetic dependencies in cancer cells lines: the DepMap project ^13,14^. We grouped cell lines according to whether or not they harbor biallelic damaging alterations in *BRCA1* or *BRCA2*, and then plotted the distribution of their gene-level depletion scores (where lower numbers indicate negative impact on cell fitness) (Table S2). Despite both datasets having only a few annotated biallelic *BRCA1-* or *BRCA2*-mutated cell lines, *CIP2A* targeting had the most penetrant, significant and profound impact on the fitness of BRCA1/2-deficient cancer cells in both datasets, with *APEX2* also showing good separation between the BRCA-proficient and -deficient groups (Fig. 1BC, S1B and Table S2). Since these studies highlighted *CIP2A* as having a particularly strong genetic interaction with *BRCA1/2*, we then used clonogenic survival assays to confirm the synthetic lethality conferred by the loss of CIP2A in the engineered *BRCA1^−/−^* and *BRCA2^−/−^* cell lines (Fig. 1D-G, S1C-E; details on sgRNA sequences and indel formation are found in Tables S3 & S4). Re-introduction of an sgRNA-resistant *CIP2A* transgene (*CIP2A\**) into *BRCA1^−/−^* and *BRCA2^−/−^* cells rescued the synthetic-lethality phenotype (Fig 1E,G). Lastly, we undertook a “reverse” CRISPR-based synthetic lethality screen with a *CIP2A* knockout query cell line (in the RPE1-hTERT *p53^−/−^* Cas9 background 15) that further confirmed synthetic lethality between CIP2A and HR genes, since *BRCA1*, *BRCA2*, *PALB2*, and *FANCM* were among the top synthetic-lethal hits, as determined by CCA and BAGEL2 16 (Fig. 1H and Table S1). We conclude that *CIP2A* is essential in a broad range of engineered and tumor-derived HR-deficient cell lines.

*CIP2A* encodes a protein of 905 amino acid residues that can be broadly split in two regions: a highly structured N-terminal half consisting of an armadillo (Arm) repeat core (residues 1-560)^17^ followed by a C-terminal half predicted to form a coiled-coil ^17^ (Fig. 1I). The exact molecular function of CIP2A is unknown although it is a reported inhibitor of the PP2A phosphatase and is overexpressed in multiple tumor types ^18,19^. Mice homozygous for a near-null *Cip2a* allele produced by gene trapping have a typical lifespan and develop normally with the exception of a mild spermatogenesis defect ^20^.

While a direct role for CIP2A in DNA repair or replication has not been reported to date, loss of CIP2A is associated with sensitivity to ATR inhibitors ^15^ and to a few other genotoxins ^21^, including the TOP1 poison camptothecin (Fig. S2A). These observations initially suggested that CIP2A may repair or prevent accumulation of replication-borne DNA lesions that require HR for their repair, since this is the basis for the *PARP1*-*BRCA* and *APEX2*-*BRCA* synthetic lethality. To test this possibility, we examined spontaneous sister-chromatid exchanges (SCEs), which are reflective of replication-associated DNA lesions that are repaired by HR ^22^. In contrast to *APEX2* sgRNAs or PARP inhibition 23, CIP2A-depleted cells experience near-basal levels of spontaneous SCEs, indicating that CIP2A loss does not greatly increase the amount of DNA lesions that engage the HR pathway (Figs. 2A and S2B). In support of this possibility, *CIP2A^−/−^* cells have similar levels of spontaneous DSBs in S phase, marked by γ-H2AX, as its parental cell line (Fig. 2B). Together, these results suggest that the *CIP2A-BRCA* synthetic lethality is not due to an increased load of replication-associated DNA lesions that are usually processed by HR.

**Fig. 2.**
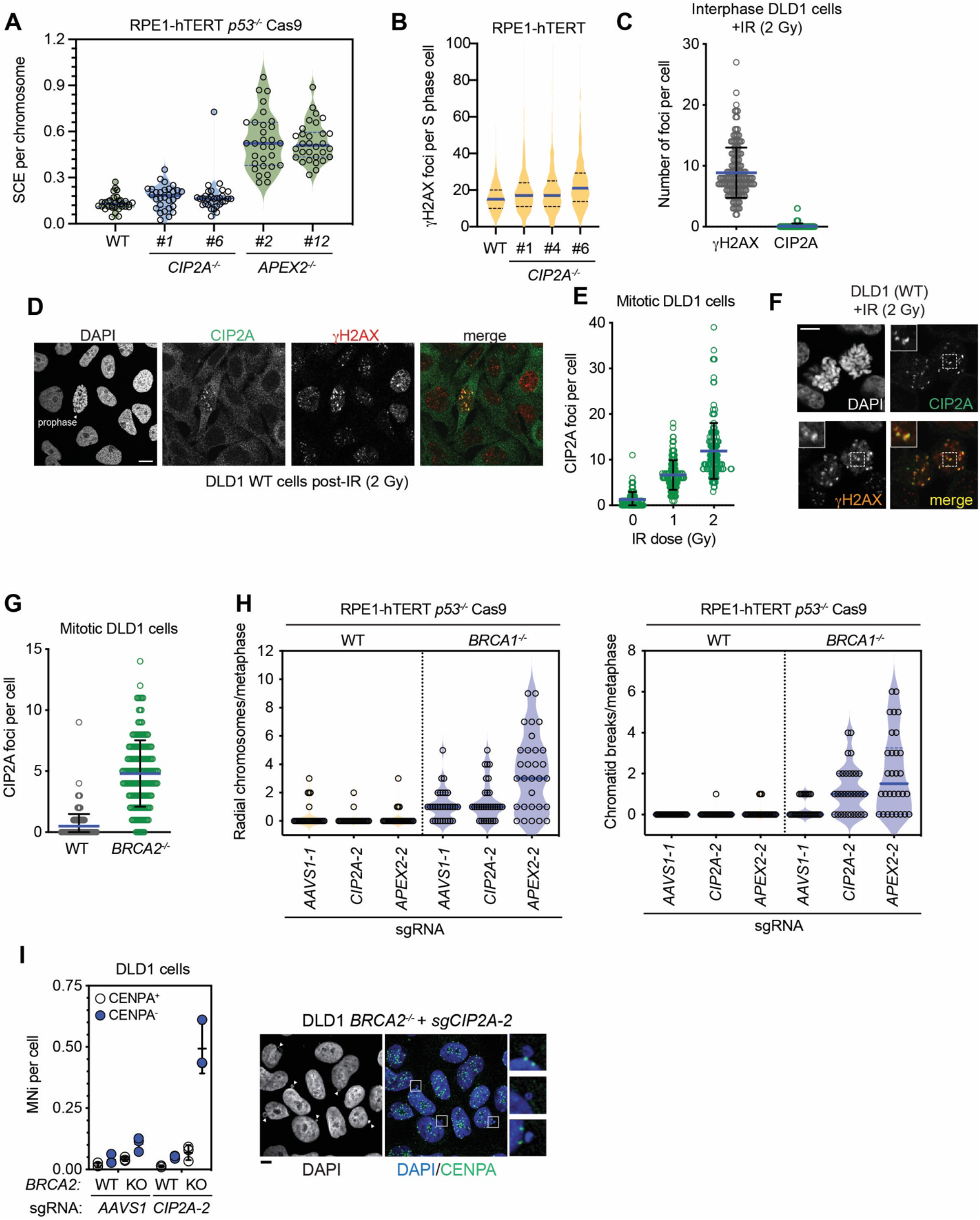
CIP2A prevents acentric chromosome segregation. (**A**) Analysis of spontaneous sister chromatid exchanges (SCEs) in RPE1-hTERT *p53^−/−^* Cas9-derived cell lines of the indicated genotype. The violin plot summarizes data from 3 biological replicates. The blue line is the median and dashed lines are at 1^st^ and 3^rd^ quartiles. (**B**) Violin plot of the automated quantitation of γH2AX foci in S phase cells of RPE1-hTERT wild-type (WT) and indicated *CIP2A^−/−^* clones. The blue line is at the median and 25^th^ and 75^th^ quartiles are indicated with dashed lines; n=8361 (WT), 3891 (*CIP2A^−/−^* #1), 2869 (*CIP2A^−/−^* #4), 1924 (*CIP2A^−/−^* #6). **(C**) Quantitation of γH2AX and CIP2A IR-induced foci, 1 h post-IR (2 Gy) in interphase DLD1 cells. Plot represents the aggregate of 3 independent experiments. The bar is at the median ± S.D. (**D**) Representative micrographs of the experiment shown in C. (**E**) Quantitation of γH2AX and CIP2A IR-induced foci, 1 h post-IR (2 Gy) in mitotic DLD1 cells. Plot represents aggregate of 3 independent experiments. The bar is at the median ± S.D. (**F**) Representative micrographs of the experiment shown in (e). Scale bar = 10 μm. (**G**) Quantitation of spontaneous CIP2A foci in mitotic DLD1 parental (WT) or *BRCA2^−/−^* cells. Plot represents aggregate of 3 independent experiments. The bar is at the median ± S.D. (**H**) Quantitation of radial chromosomes (left) and chromatid breaks (right) in metaphase spreads from RPE1-hTERT *p53^−/−^* Cas9 cells upon transduction of virus expressing sgRNAs targeting *AAVS1*, *APEX2* or *CIP2A* (10 metaphases scored from at least 2 biologically independent experiments). Representative images are shown in Fig. S2E. (**I**) Quantitation of micronuclei (MNi) staining positive (+) or negative (−) for CENPA in DLD1 cells, parental (WT) or *BRCA2^−/−^* (KO), 7 d post-transduction with indicated sgRNAs. Biological replicates are shown and the bars represent the mean ± S.D. Representative micrographs are shown on the right. Arrowheads point at micronuclei. Scale bar = 10 μm.

## CIP2A acts on mitotic DNA lesions

A lack of direct involvement of CIP2A in DNA repair or DNA replication is further supported by the subcellular localization of CIP2A. As previously noted ^24^, CIP2A is cytoplasmic in interphase cells as determined by immunofluorescence microscopy (Fig. S2C). DNA damage caused by ionizing radiation (IR) did not promote CIP2A translocation from the cytoplasm to the nucleoplasm in interphase cells, but rather led to a striking formation of IR-induced, γH2AX-colocalising CIP2A foci solely in mitotic cells (Fig. 2C-F). We also observed an increased frequency of mitotic CIP2A foci in *BRCA2^−/−^* cells over their wild-type counterparts (Figs. 2G and S2D), suggesting that CIP2A responds to DNA damage only during M phase and that this response is likely relevant to the *CIP2A-BRCA* synthetic lethality.

The metaphases of HR-deficient cells treated with PARP inhibitors or depleted of APE2 display increased numbers of radial chromosomes ^6,25^, which are likely to be caused by the unscheduled action of non-homologous end-joining on DNA lesions that are normally repaired by HR. Depletion of CIP2A in *BRCA1*^−/−^ cells did not increase radial chromosome formation but we did detect an increase in chromatid breaks (Figs. 2H and S2E). Together with the near-normal SCE frequency of *CIP2A^−/−^* cells, these results further indicate that CIP2A must support the viability of HR-deficient cells via a mechanism distinct from PARP or APE2. A clue to this mechanism emerged when we observed that depletion of CIP2A led to a striking increase in the frequency of micronuclei (MNi) formed in *BRCA2*^−/−^ cells (Figs. 2I and S2F). These micronuclei were largely CENPA-negative, indicating that they originate from the mis-segregation of acentric (i.e. broken) chromosomes (Fig. 2I). These results suggest that CIP2A promotes the viability of HR-deficient cells by guarding against the formation and/or mis-segregation of acentric chromosomes.

## A CIP2A-TOPBP1 complex

The micronucleation caused by loss of CIP2A in HR-deficient cells was revealing in light of work showing that MDC1 and TOPBP1 form mitotic IR-induced foci that promote segregation of broken chromosomes ^26^. This was not the sole connection to these genes since analysis of DepMap data ^13^ showed essentiality profiles for *MDC1* and *TOPBP1* that are highly correlated to those of *CIP2A* (Fig. 3A). Similarly, genotoxin sensitivity profiles generated from a DNA damage chemogenomic dataset 21 also links *CIP2A* to *MDC1* (Fig. S3A). Together, these data hinted that CIP2A collaborates with MDC1 and TOPBP1 to promote the accurate segregation of damaged chromosomes. In support of this possibility, CIP2A, MDC1 and TOPBP1 colocalized at IR-induced mitotic foci in nocodazole-treated cells (Fig. 3B). Protein depletion studies with siRNAs further showed that MDC1 was acting upstream of TOPBP1 and CIP2A, and that the localization of TOPBP1 and CIP2A to mitotic broken chromosomes was dependent on each other (Figs. 3C and S3C).

**Fig. 3.**
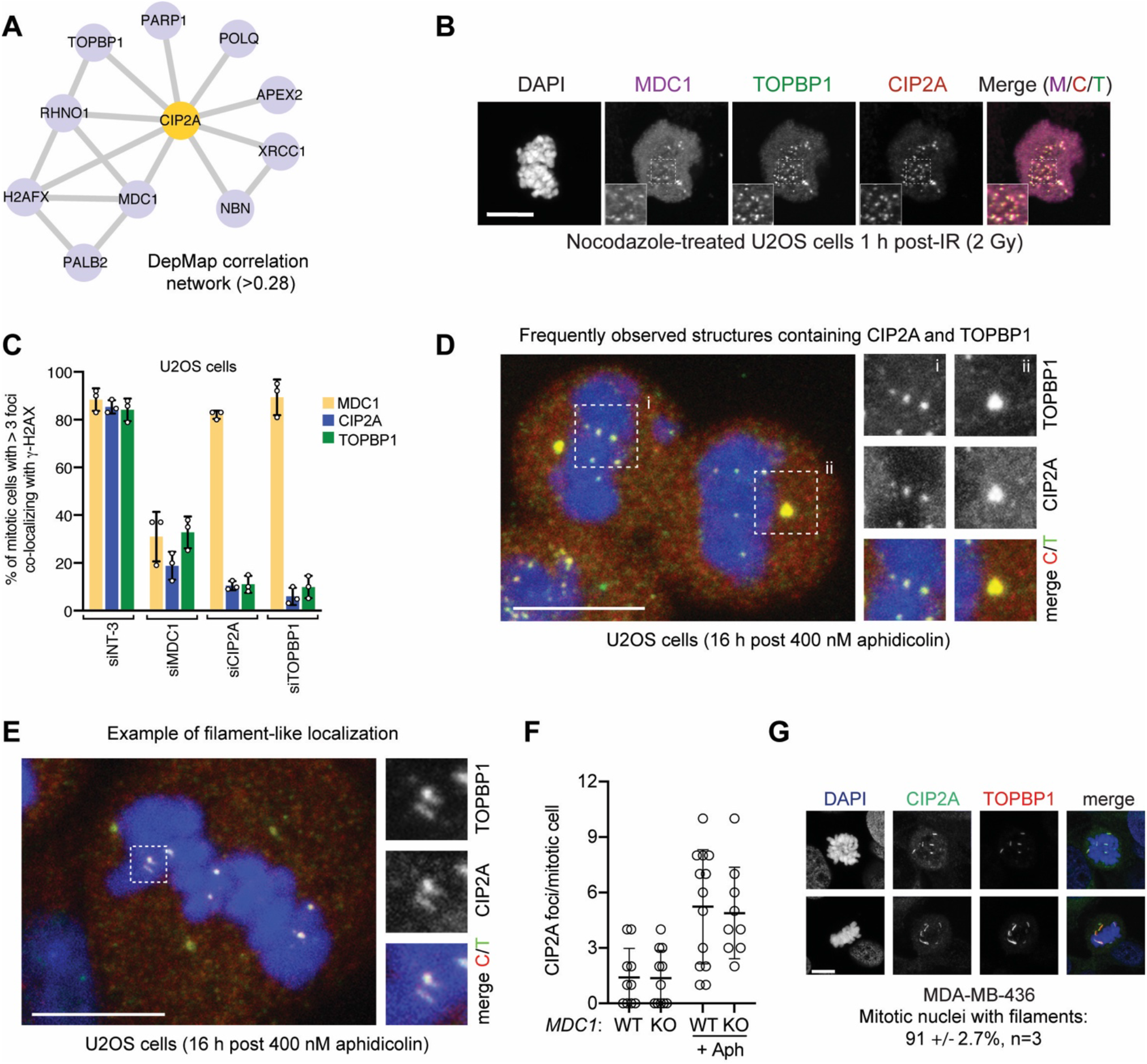
CIP2A co-localizes with TOPBP1 on mitotic structures. (**A**) Correlation network based on Pearson correlation of gene-level dependency scores (>0.28) derived from the Broad DepMap data. (**B**) Representative micrograph of an X-irradiated (2 Gy) mitotic U2OS cell treated with 100 ng/mL nocodozole for 16 h and stained with the indicated antibodies. DNA was stained with DAPI. Scale bar = 10 μm. (**C**) Quantitation of MDC1, CIP2A and TOPBP1 IR-induced foci in mitotic U2OS cells treated with nocodazole and the indicated siRNAs (siNT-3 is a non-targeting control). Data is presented as the mean ± S.D. (n=3). Representative micrographs are shown in Fig. S3B. (**D, E**) Types of CIP2A/TOPBP1 structures observed in mitotic cells after treatment with low dose aphidicolin. Maximum intensity projections of confocal z-stacks of U2OS wild type mitotic cells treated with 400 nM aphidicolin for 16 h. Scale bars = 10 μm. Besides centrosomes (D, inset *ii*) that always stain for TOPBP1 and CIP2A regardless of the treatment, small round foci are the most frequently observed structures in response to aphidicolin treatment (D, inset *i*). Less frequently observed structures include curved and straight filamentous assemblies (E and Fig. S3D). (**F**) Quantitation of CIP2A and TOPBP1 colocalizing foci in U2OS (WT) and *MDC1^−/−^* (KO) cells after treatment with 400 nM aphidicolin (16 h). The number of foci per mitotic cell are shown and the bars represent the mean ± S.D. (**G**) Representative micrographs of MDA-MB-436 mitotic cells stained for CIP2A and TOPBP1. DNA was stained with DAPI. Scale bars = 10 μm. Quantitation of the percentage of cells with filaments is indicated.

While the above data suggest that CIP2A acts downstream of MDC1 in promoting the segregation of MDC1-marked broken chromosomes, it also raised a conundrum since loss of MDC1 does not cause lethality in BRCA2-deficient cells (Fig. S3B). This observation indicates that the MDC1-dependent modulation of DSBs in mitosis is not relevant to the CIP2A-BRCA synthetic lethality.

However, TOPBP1 is known to have MDC1-independent roles in promoting genome integrity during M phase as it also responds to the presence of incompletely replicated DNA that persists until mitosis 26-28. Analysis of TOPBP1 and CIP2A localization on mitotic chromosomes of wild type or *MDC1*^−/−^ U2OS cells showed that TOPBP1 and CIP2A colocalized in a number of structures in the absence of IR exposure, and that the frequency of these structures was stimulated by low-dose treatment (400 nM) with aphidicolin, a DNA polymerase inhibitor (Figs. 3D-F and S3D). The aphidicolin-stimulated structures included small foci that are often symmetrically distributed between the dividing chromatin masses (Fig. 3D, inset *i*) as well as filament-like structures that most often occur within the chromatin of the dividing daughter cells, distinguishing them from ultrafine bridges (UFBs) ^29^ (Figs. 3E and S3D). We also observed that CIP2A and TOPBP1 localized to centrosomes, a known site of TOPBP1 and CIP2A localization ^30,31^ (Fig. 3D, inset *ii*). Centrosomal localization is seen in every cell irrespective of treatment whereas the CIP2A-TOPBP1 foci and filaments were rare in untreated HR-proficient cells, but their frequency could be increased by aphidicolin treatment in a manner that was mostly independent of MDC1 (Figs. 3F and S3E). Remarkably, in the tumor-derived cell line MDA-MB-436, which is defective in BRCA1^32^, CIP2A-TOPBP1 filaments were present in nearly all mitotic cells examined (91 ± 2.7%, n=3; Figs. 3G and S3F). In MDA-MB-436 cells, the CIP2A-TOPBP1 filaments appear to be seeded from chromosomal loci in mitosis but seem to elongate over time and could sometimes be observed as detached from the chromatin mass in some dividing cells (Fig. S3F). While the nature of these intriguing structures remains under investigation, our data suggest they are initially formed as a consequence of unresolved replication-associated DNA lesions and are thus likely relevant to the *CIP2A-BRCA* synthetic lethality.

The intimate and interdependent localization of CIP2A and TOPBP1 on mitotic structures hinted that they may interact with each other. Indeed, CIP2A retrieves TOPBP1 in co-immunoprecipitation assays (Fig. 4A). The two proteins were also found to interact in a cellular co-localization assay where TOPBP1 fused to the LacR DNA-binding domain is targeted to a chromosomal site with ~256 copies of the LacO sequence integrated (Fig. 4B-D). The LacR/LacO assay was conducted with interphase cells, suggesting that some CIP2A can shuttle in and out of the nucleus. The TOPBP1-binding region on CIP2A mapped to the highly structured Arm-repeat core (residues 1-560; Fig. 4D), a finding we confirmed by yeast two-hybrid analysis, which also suggested that the interaction is direct (Fig. 4E).

**Fig. 4.**
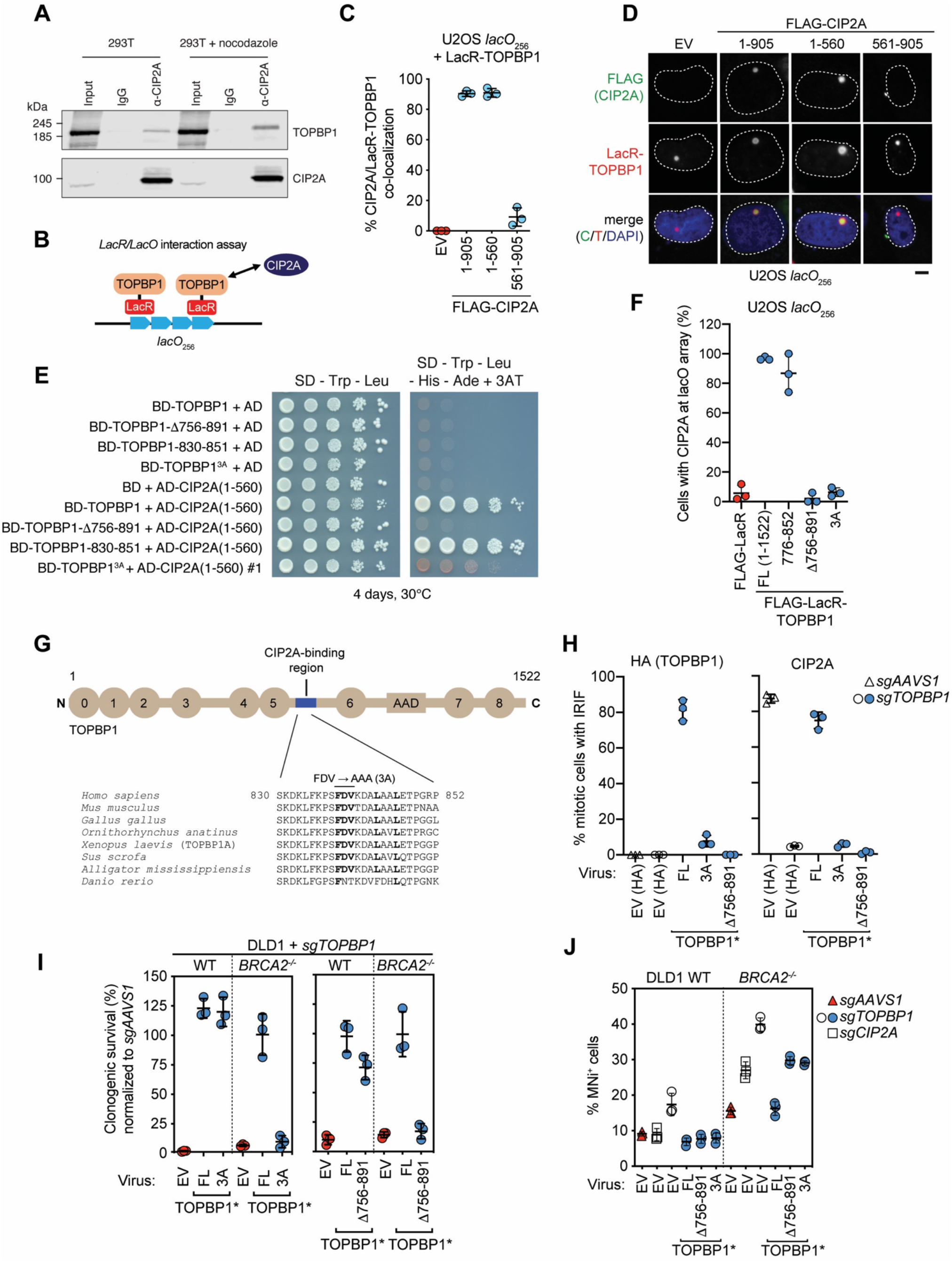
The CIP2A-TOPBP1 interaction is essential in *BRCA2^−/−^* cells. (**A**) Co-immunoprecipitation of CIP2A with TOPBP1. Whole-cell extracts from 293T cells, untreated or treated with nocodazole for 16 h, were subjected to immunoprecipitation with normal mouse IgG or a CIP2A antibody and were then immunoblotted with TOPBP1 (top) or CIP2A (bottom) antibodies. (**B**) Schematic of the LacR/LacO assay. (**C**, **D**) LacR/LacO assay assessing the interaction of Flag-tagged CIP2A and deletion mutants with LacR-TOPBP1 in U2OS *lacO*_*256*_ cells. Quantitation of the assay is in C where 3 biological replicates are shown and the bars represent the mean ± S.D. Representative micrographs are shown in D. Scale bar = 10 μm. (**E**) Yeast two-hybrid assay for interaction between TOPBP1 variants and CIP2A (1-560). Expression of proteins was verified by immunblotting but not shown. (**F**) LacR/LacO assay assessing the interaction between endogenous CIP2A and TOPBP1 variants fused to Flag-LacR. Data points represent biological replicates and data is presented as the mean ± S.D. FL=full-length. (**G**) Schematic of TOPBP1 and sequence conservation of the minimal CIP2A-interaction motif. (**H**) Quantitation of CIP2A and HA-tagged TOPBP1 mitotic foci in DLD1 cells stably expressing full-length (FL) or the indicated mutants of sgRNA-resistant *TOPBP1* (TOPBP1*) or empty virus encoding only the HA tag (EV(HA)) followed by transduction of viruses expressing both Cas9 and sgRNAs targeting *TOPBP1* (*sgTOPBP1*) or AAVS1 (*sgAAVS1*). Data points represent biological replicates and the bars represent the mean ± S.D. (n=3). (**I**) Clonogenic survival of DLD1 wild-type (WT) and *BRCA2^−/−^* cells stably expressing sgRNA-resistant *TOPBP1* (TOPBP1*, FL), the indicated *TOPBP1* mutants, or an empty virus (EV) followed by inactivation of the chromosomal copies of TOPBP1 with an sgRNA and Cas9 (*sgTOPBP1*). Quantitation of the data is shown in I where representative images of the crystal violet-stained colonies are shown Fig. S4F. Data points represent biological replicates, and the error bars represent the mean ± S.D. n=3. (**J**) Quantitation of micronuclei (MNi) in DLD1 wild-type (WT) and *BRCA2^−/−^* cells stably expressing sgRNA-resistant *TOPBP1* (TOPBP1*), the indicated *TOPBP1* mutants, or an empty virus (EV) followed by inactivation of *TOPBP1*, *CIP2A* or *AAVS1* with the indicated sgRNAs and Cas9. Data points represent biological replicates, and the bars represent the mean ± S.D. n=3.

We mapped the CIP2A-interacting region of TOPBP1 to a region located between BRCT5 and BRCT6, in a segment encompassing residues 830-851 (by yeast two-hybrid; Fig. 4e) or residues 776-852 (with LacR/LacO; Figs. 4F and S4AB). Deletion of a segment of TOPBP1 comprising this region, i.e. TOPBP1-Δ756-891, completely abolished the CIP2A-TOPBP1 interaction as monitored by the LacR/LacO system or yeast two-hybrid (Fig. 4EF). We then used yeast two-hybrid analysis to identify five point mutants that disrupt binding between CIP2A and TOPBP1 (Fig. S4C), which included mutations that targeted a conserved three-residue segment on TOPBP1, F837-D838-V839 (Fig. 4G). Mutation of these residues to alanine in the context of full-length TOPBP1 generated the TOPBP1^3A^ mutant, which has impaired interaction with CIP2A in both yeast and mammalian cells (Fig. 4EF).

The identification of TOPBP1 variants defective in CIP2A binding enabled us to test whether the TOPBP1-CIP2A interaction was essential for the viability of BRCA-deficient cells. We generated DLD1 *BRCA2^−/−^* cell lines stably transduced with sgRNA-resistant lentiviruses that express TOPBP1, TOPBP1^3A^ and TOPBP1-Δ756-891 (Fig. S4D) and then inactivated the endogenous chromosomal copies of *TOPBP1* by Cas9-mediated mutagenesis. As hinted by the depletion studies, cells expressing TOPBP1-Δ756-891 and TOPBP1^3A^ failed to form mitotic CIP2A IR-induced foci (Figs. 4H and S4E) and displayed rapid loss of fitness selectively in the *BRCA2^−/−^* background upon removal of endogenous *TOPBP1* (Figs. 4I and S4F). The lethality of TOPBP1-Δ756-891 and TOPBP1^3A^ in *BRCA2^−/−^* cells was also accompanied with an increase in micronucleation, suggesting lethal chromosome instability (Figs. 4J and S4G). We conclude that the CIP2A-TOPBP1 interaction is essential for the viability of HR-deficient cells.

## Therapeutic proof-of-concept

Our data suggest that disrupting the CIP2A-TOPBP1 interaction may be an attractive therapeutic strategy. To model this approach, we identified a fragment of TOPBP1 corresponding to residues 756-1000 (Fig. 5A), referred to as “B6L” (for BRCT6-long) that is highly effective at disrupting mitotic CIP2A foci when expressed from a lentiviral vector (Figs. 5B and S5A). B6L expression is under the control of a FKBP12-derived destabilization domain (DD) ^33^, which enables tight induction of B6L expression upon addition of the Shield-1 or water-soluble AS-1 (Aqua-Shield-1) compounds (Fig. S5B). Incucyte imaging of *BRCA2^−/−^* cells following induction of B6L showed a near-complete cessation of proliferation in DLD1 *BRCA2^−/−^* cells within 3 days of induction whereas it was innocuous to its BRCA-proficient parental cell line (Fig. 5CD). We also observed that induction of B6L for 2 days followed by a washout of AS1 (Fig. 5E) led to an irreversible cessation of growth as determined by clonogenic survival (Figs. 5F and S5C) and was accompanied by rapid and high levels micronucleation, further suggesting that segregation of acentric fragments is a plausible cause of cell death in BRCA-deficient cells (Fig. 5G). Disruption of the CIP2A-TOPBP1 interaction with B6L did not impair ATR signaling, ruling out that the impact of B6L is due to ATR misregulation (S5D). B6L expression also caused micronucleation and impaired proliferation of the tumor-derived MDA-MB-436 cell line, indicating its ability to stunt proliferation of BRCA-deficient cells and cause chromosome mis-segregation is not limited to engineered backgrounds (Figs. 5G-I and S5EF).

**Fig. 5.**
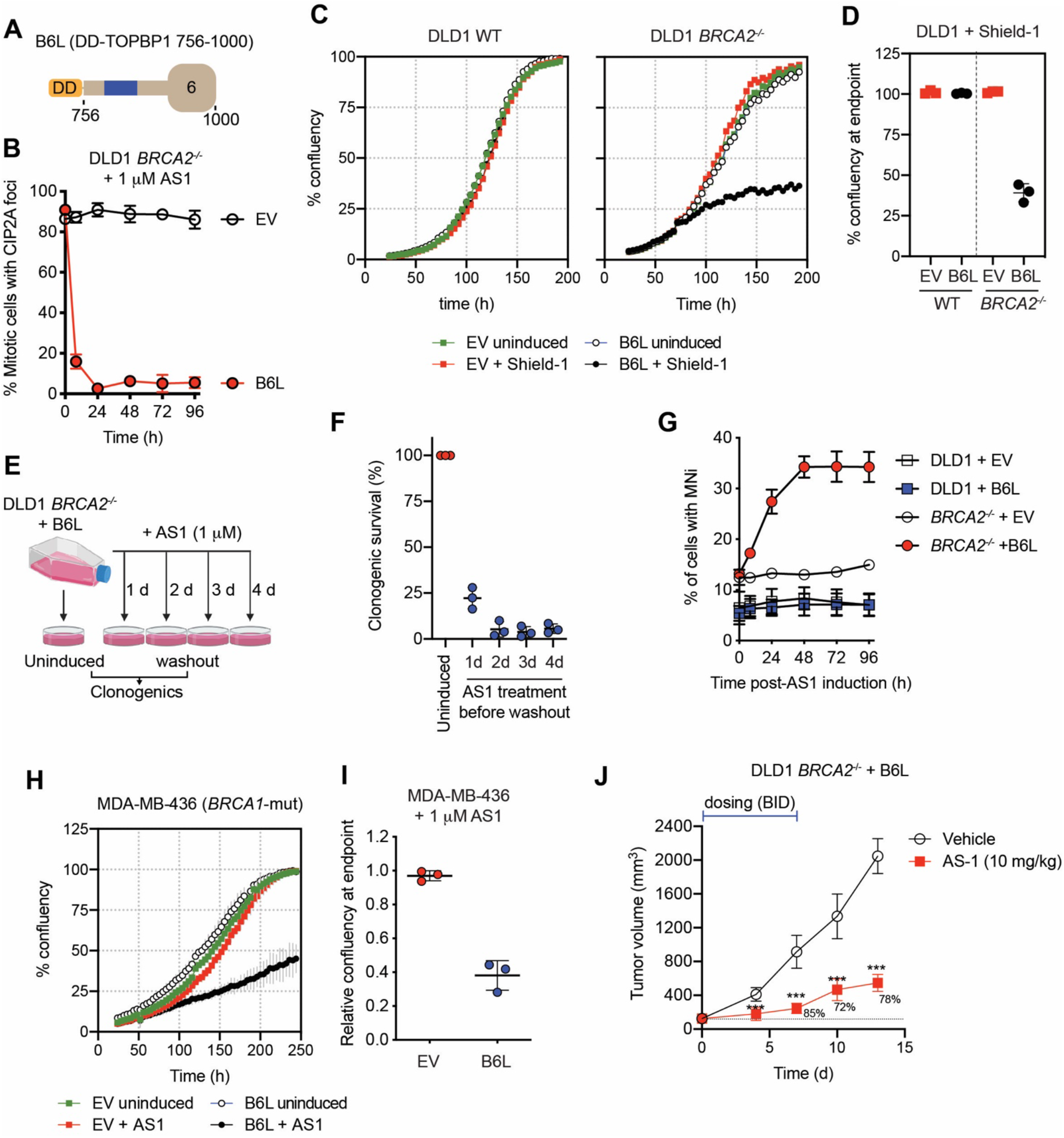
Therapeutic proof-of-concept. (**A**) Schematic of B6L, a fragment derived from TOPBP1 residues 756-1000 fused to the destabilization domain (DD). (**B**) Quantitation of mitotic CIP2A foci in DLD1 *BRCA2^−/−^* upon B6L stabilization. Data is shown as mean ± S.D. (n=3). (**C**) Representative proliferation curves for DLD1 parental (left) and *BRCA2^−/−^* (right) cells upon B6L stabilization by Shield-1 treatment (1 μM). Cells were transduced with an empty virus (EV) that expresses the DD domain as control. (**D**) Aggregate of 3 biological replicates of the experiment shown in (c). Data is presented as mean ± S.D. (n=3). (**E**) Schematic of the experiment shown in F. (**F**) Clonogenic survival of DLD1 *BRCA2^−/−^* cells following expression of B6L for the indicated periods of time. Data is presented as mean ± S.D. (n=3). (**G**) Quantitation of micronuclei (MNi)-positive cells in DLD1 WT or *BRCA2^−/−^* cells following addition of AS1. Data presented as mean ± S.D. (n=3). (**H**) Representative proliferation curves for MDA-MB-436 cells upon B6L stabilization by AS1 treatment (1 μM). Cells were transduced with an empty virus (EV) that expresses the DD domain as control. (**I**) Aggregate of 3 biological replicates of the experiment shown in H. Data presented as the mean ± S.D. (**J**) Growth of tumor xenografts derived from DLD *BRCA2*^−/−^ cells transduced with a B6L-encoding lentivirus treated intraperitoneally BID for 7 d either with AS-1 (10 mg/kg) or with vehicle. After termination of treatment, tumors were grown and monitored without for an additional 8 d. Data is presented at the mean ± S.D. (n=8). Tumor growth inhibition is indicated, and significance (*** p<0.001) was determined by a two-tailed unpaired Student’s t-test. See Fig. S6 for additional controls and pharmacokinetics of the AS-1 compound.

Finally, to test whether disrupting the CIP2A-TOPBP1 interaction could inhibit tumor growth, we established tumors from DLD1 *BRCA2^−/−^* cells transduced either with empty (EV) or B6L-expressing lentivirus in NOD-SCID mice. We first characterized the pharmacokinetic properties of the AS-1 compound and found it to be poorly bio-available, but we could optimise a dosing regimen (intraperitoneal injection) that resulted in plasma concentrations that could exceed the EC50 concentration necessary to inhibit growth *BRCA2^−/−^* cells for up to 4 hrs per day when administered twice daily (BID dosing; Fig. S6AB). Despite being a suboptimal dosing regimen, the periodic stabilization of B6L in *BRCA2^−/−^* tumors was sufficient to cause striking tumor growth inhibition over the course of a 7-day treatment, reaching 85% (Fig. 5J). Furthermore, tumor growth inhibition was maintained until the completion of the experiment, 8 days after administration of the last dose of AS-1 (Fig. 5J). We conclude that not only is the inhibition of the CIP2A-TOPBP1 interaction providing an attractive therapeutic strategy for HR-deficient cancers but our findings indicate that the complete and sustained inhibition of the CIP2A-TOPBP1 interaction may not be necessary for achieving BRCA-deficient tumor control.

## Discussion

The observation that acute inactivation of *BRCA1* and *BRCA2* causes cellular lethality is in line with a model where BRCA1/2-deficient tumors acquire genetic and/or non-genetic adaptive mechanisms that enable these cells to proliferate in the face of HR deficiency. While p53 inactivation is one genetic means by which cells acquire the ability to tolerate HR-deficiency ^34,35^, our findings suggest that a dependency on the CIP2A-TOPBP1 complex provides another way to endow HR-deficient cells with the ability to proliferate. Our observations strongly suggest a role for this complex in promoting the segregation of chromosomes that did not fully complete DNA replication (Fig. S6D). We speculate that CIP2A either participates in the resolution of incompletely replicated chromosomes or, more likely, that it participates in physically bridging acentric fragments to their centromere-bearing counterpart following the mitotic processing of incompletely replicated chromosomes (Fig. 5J). While the molecular details of this function remain to be delineated, we anticipate that this role of CIP2A-TOPBP1 will be distinct from other mitotic DNA damage tolerance pathways ^29^ since the genes encoding proteins known to have central roles in these processes, such as RAD52 (MiDAS) or PICH (ultrafine bridge resolution), were not synthetic-lethal with *BRCA1/2* in either of the CRISPR screens we undertook (Table S1).

In conclusion, we nominate the CIP2A-TOPBP1 interaction as a therapeutic target for the treatment of HR-deficient tumors. Since the loss of CIP2A does not cause high loads of DNA damage in HR-proficient cells, and since *Cip2a*-deficient mice develop normally with a typical lifespan, we predict that inhibiting the CIP2A-TOPBP1 interaction will have non-overlapping toxicity with PARP inhibitors, and thus could enable a greater range of therapeutic combinations. Furthermore, since we observed that CIP2A loss impairs fitness in a model of PARP inhibitor resistance, *BRCA1^−/−^ 53BP1^−/−^* cells (Fig. S6E), inhibitors of the CIP2A-TOPBP1 interaction may also prove effective in subsets of patients that progress on PARP inhibitor therapy. Efforts to discover small molecule inhibitors of the CIP2A-TOPBP1 interaction are currently underway.

## Acknowledgments

We thank members of the Durocher lab for helpful discussions and Alain Jeanrenaud for technical help. We also thank Jason Moffat for his generous sharing of the TKO sgRNA libraries. We are grateful to Kin Chan at the NBCC (LTRI) for sequencing. We are also grateful to the Repare in vivo pharmacology team (Anne Roulston, Alexanne Bonneau-Fortin, Marie-Ève Leclaire and Sara Fournier) for their help with tumor xenograft assays. SA was supported by a Banting post-doctoral fellowship and SER was supported by a fellowship from AIRC. AAQ holds and DS held postdoctoral fellowships from the Canadian Institutes for Health Research (CIHR). DD is a Canada Research Chair (Tier I) and work in the DD lab was supported by grants from the CIHR (FDN143343) and Canadian Cancer Society (705644) with additional support from the Krembil Foundation and Repare Therapeutics.

## Data availability statement

All data generated or analysed during this study are included in this published article (and its supplementary information files).

## Code availability

Details about CRISPRcount Analysis (CCA) can be found in the methods, including instructions on how to install the software. Code is also available on github at https://github.com/tohsumi-repare/cca

## METHODS

### Cell culture

RPE1-hTERT, U2OS and 293T cells were grown at 37°C and 5% CO_2_ in DMEM supplemented with 10% FBS (Wisent #080150) and 1% Pen/Strep (Wisent). Parental and *BRCA2^−/−^* DLD1 cells were purchased from Horizon and maintained in RPMI-1640 medium (ATCC 30-2001) supplemented with 10% FBS and 1% Pen/Strep. Parental and *BRCA2^−/−^* DLD1 Cas9 cells were generated through viral infection with lentiCas9-Blast (Addgene #52962) followed by blasticidin selection. MDA-MB-436 cells were purchased from ATCC and maintained in DMEM supplemented with 10% FBS and 1% Pen/Strep. DLD1 and MDA-MB-436 cell lines were grown at 37°C in a low-oxygen (3% O_2_) incubator. The RPE1-hTERT *p53^−/−^ BRCA1^−/−^*, *BRCA1^−/−^53BP1^−/−^*, *APEX2^−/−^*-*, CIP2A^−/−^* and the U2OS *MDC1^−/−^* knockout cell lines were described previously^6,15,26,36^. The *CIP2A^−/−^* RPE1-hTERT cell line (i.e. p53^+^) is described in de Marco Zompit et al. (submitted).

### Lentiviral transduction

Lentiviral particles were produced in 293T cells in 10-cm plates by co-transfection of 10 μg of targeting vector with 3 μg VSV-G, 5 μg pMDLg/RRE and 2.5 μg pRSV-REV (Addgene #14888, #12251, #12253) using calcium phosphate. Viral transductions were performed in the presence of 4 μg/μL polybrene (Sigma-Aldrich) at a multiplicity of infection (MOI) <1. Transduced cells were selected by culturing in the presence of blasticidin (InvivoGen) or nourseothricin (Jena Bioscience) depending on the lentiviral vector used.

### Two-color competitive growth assays

Cells were transduced with sgRNA expression lentiviruses, either expressing NLS-mCherry-sg*AAVS1* (control) or an NLS-GFP-sgRNA targeting a specific gene of interest (Supplementary Table 3) at an MOI of ~0.5. 24 h after transduction, cells were selected for 48 h using 15 μg/mL (RPE1) or 2 μg/mL (DLD1) puromycin (Life Technologies). 96 h after transduction, mCherry- and GFP-expressing cells were mixed 1:1 (2,000 cells each for RPE1; 3,000 cells each for RPE1 *BRCA1^−/−^* and DLD1; 9,000 cells each for DLD1 *BRCA2^−/−^*) and seeded in a 12-well plate. Cells were imaged for GFP and mCherry 24 h after initial plating (t=0) and at the indicated timepoints using a 4X objective InCell Analyzer system (GE Healthcare Life Sciences, Marlborough). Segmentation and counting of GFP- and mCherry-positive cells were performed using an Acapella script (PerkinElmer, Waltham). Efficiency of indel formation was analysed by performing PCR amplification of the region surrounding the sgRNA sequence using DNA isolated from cells collected from 4 to 7 days after transduction and subsequent ICE analysis (https://ice.synthego.com/#/) (Supplementary Table 4).

### Clonogenic survival assays

Cells were transduced at low MOI (<1.0) with lentivirus derived from pLentiGuide (RPE1 cells) or pLentiCRISPRv2, which expressed sgRNAs targeting *CIP2A, TOPBP1* or *AAVS1* (which was used as control). Puromycin-containing medium was added the next day to select for transductants and cells were seeded for clonal growth 48 h later. Cells were seeded in 10-cm dishes (750-5,000 cells per 10 cm plate, depending on cell line and genotype). For drug sensitivity assays, cells were seeded into media containing a range of camptothecin (Sigma) concentrations (for determination of camptothecin sensitivity) or in regular media after several days of AS1 treatment (i.e. after induction of B6L). For clonogenic survival assays performed with *CIP2A^−/−^* cells, plates were incubated in atmospheric oxygen. Experiments performed with *BRCA1^−/−^* and *BRCA2^−/−^* cells and their controls were incubated at 3% O_2_. Medium was refreshed after 7 d. After 14-20 d, colonies were stained with a crystal violet solution (0.4% (w/v) crystal violet (Sigma), 20% methanol). Colonies were manually counted or counted using a GelCount instrument (Oxford Optronix). Data were plotted as surviving fractions relative to untreated cells or *sgAAVS1*-transduced controls.

### Plasmids

For CRISPR-Cas9 genome editing, sgRNAs were cloned either in lentiCRISPRv2 or in lentiguide-NLS-GFP as in ref^6^. The sgRNA sequences used in this study are included in Supplementary Table 3. The pcDNA5-FRT/TO-LacR-FLAG-TOPBP1 plasmid was obtained from Addgene (#31313). Point mutants were introduced by site-directed mutagenesis using Quikchange (Agilent). For TOPBP1 rescue experiments, the pLenti-CMVie-IRES-BlastR (pCIB) plasmid was obtained from Addgene (#119863). pCIB-2xHA was generated by cloning a double HA tag with a flanking NotI site in pCIB, using AscI and BamHI restriction sites. The *TOPBP1* coding sequence was amplified from pcDNA5-FRT/TO-LacR-FLAG-TopBP1 and mutagenised at Genscript (Piscataway, NJ) to generate an sgRNA-resistant construct with a silent mutation at Thr263 (ACC to ACA). This fragment was then cloned into pCIB-2xHA using NotI and BamHI restriction sites to generate pCIB-2xHA-TOPBP1-sgR. For the inducible expression of the B6L fragment, we first synthesized a cassette coding the FKBP-derived destabilization domain (DD)^33^ along with an EcoRI restriction site and a single FLAG tag (Genscript). This cassette was then cloned into pHIV-NAT-T2A-hCD52 (kind gift of R. Scully) using the NotI and BamHI restriction sites to generate pHIV-NAT-DD-FLAG. pHIV-NAT-DD-FLAG-B6L was amplified by PCR from pcDNA5-FRT/TO-LacR-FLAG-TOPBP1(756-1000) and cloned into pHIV-NAT-DD-FLAG using EcoRI and BamHI sites. The *CIP2A* coding sequence was amplified from a BirA-CIP2A expression plasmid (a kind gift from A.-C. Gingras) and cloned into the pcDNA5-FRT/TO-FLAG vector using the AscI and EcoRI sites. The mutation making this vector resistant to *sgCIP2A-2* (silent mutation in Ala650, GCC to GCA) was introduced by site-directed mutagenesis generating pcDNA5-FRT/TO-Flag-CIP2A-sg2R. Using this vector as a template, FLAG-CIP2A or portions of *CIP2A* were amplified by PCR and cloned into the pHIV-NAT-T2A-hCD52 using NotI and EcoRI restriction sites. The corresponding control vector, pHIV-NAT-FLAG-T2A-hCD52, and pHIV-NAT-FLAG-CIP2A(560-915) were generated by deletion PCR from pHIV-NAT-FLAG-CIP2A-sg2R. For yeast two hybrid experiments, a fragment corresponding to CIP2A (1-560) was cloned by Genscript into pGADT7 AD (Clontech/Takara) to create a fusion with the GAL4 activating domain using EcoRI and XmaI restriction sites, whereas a TOPBP1 fragment corresponding to residues 2-1523 was amplified from pCDNA5-FRT/TO-LacR-FLAG-TopBP1 and cloned into pGBKT7 (Clontech/Takara) to create a fusion with the GAL4 DNA binding domain using the NdeI and XmaI sites. pGBKT7-GAL4-BD-TOPBP1-Δ756-891 and pGBKT7-GAL4-BD-TOPBP1-3A were derived from pGBKT7-GAL4-BD-TOPBP1, removing the sequence coding residues 756-891 by deletion PCR and mutating the codons for Phe837, Asp838, Val839 to Ala by Quikchange site-directed mutagenesis, respectively. pGBKT7-GAL4-BD-TOPBP1(830-851) was generated by cloning a *TOPBP1* fragment corresponding to residues 830-851 into pGBKT7 using the NdeI and XmaI restriction sites. The alanine scanning library of the TOPBP1 830-851 fragment was generated at Genscript and cloned into pGBKT7-GAL4-BD.

### CRISPR screens

The CRISPR screens were carried out using protocols derived from refs^6,9,37^. Synthetic lethality screens are basically undertaken as two parallel screens with a parental cell line and an isogenic variant with one genetic alteration, in this case *BRCA1* or *BRCA2* loss-of-function mutations. For the BRCA2 screen, DLD1 parental and *BRCA2−/−* cells were transduced with the lentiviral TKOv3 sgRNA library 37,38 at a low MOI (~0.3) and media containing puromycin (Life Technologies) was added the next day to select for transductants. The following day, cells were trypsinized and replated in the same plates while maintaining puromycin selection. 3 d after infection, which was considered the initial time point (t0), cells were pooled together and divided into 2 sets of technical replicates. Cells were grown for a period of 18-30 d and cell pellets were collected every 3d. Each screen was performed as a technical duplicate with a theoretical library coverage of ≥ 400 cells per sgRNA maintained at every step. Genomic DNA was isolated using the QIAamp Blood Maxi Kit (Qiagen) and genome-integrated sgRNA sequences were amplified by PCR using NEBNext Ultra II Q5 Master Mix (New England Biolabs). i5 and i7 multiplexing barcodes were added in a second round of PCR and final gel-purified products were sequenced on an Illumina NextSeq500 system at the LTRI NBCC facility (https://nbcc.lunenfeld.ca/) to determine sgRNA representation in each sample.

### CRISPRCount Analysis (CCA)

CCA is a scoring approach optimized for isogenic CRISPR screens that provides gene-level scores and ranking of genes according to the impact of their targeting sgRNAs between test and control samples. CCA also aims to prioritize sgRNAs that are selectively deleterious to fitness in the test samples. CCA is available on Docker. To download the Docker image of CCA, install Docker (https://www.docker.com/) and then in a terminal window, execute: “docker pull tohsumirepare/cca”. The CCA Docker image source is located at https://github.com/tohsumi-repare/cca and the documentation for CCA, such the input file format and method of execution, is in the doc folder.

CCA employs non-parametric statistics. Implementation of CCA was based on MolBioLib^39^ (sourceforge.net/projects/molbiolib) and includes the Mann-Whitney U test from ALGLIB C++ (www.alglib.net). The input of CCA is a matrix of samples versus sgRNAs where the entries are the sgRNA readcounts in that sample. The CCA score is computed as follows: (1) normalization of the readcount file so that each sample’s count over all sgRNAs is 10 million; (2) removal of sgRNAs with readcounts at T0 is <30 to avoid false positives due to low readcounts; (3) computing a depletion matrix of samples versus sgRNAs where the depletion = 1 – (count at final time)/(count at initial time) = 1 – foldchange. The depletion is such that it is maximum, 1, if the test sample has no viable cells at the final timepoint. The depletion may be negative if there is proliferation of cells at the final timepoint. By default, we limit the minimum value of the depletion of all control samples to 0 (doing otherwise can create false positive hits if sgRNAs cause proliferation in control samples).

For a given gene, we let the vector of depletion values over all test samples be denoted t and over all control samples be denoted c. For vector v, let Q3(v) be the third quantile of v. The CCA score for that gene is:

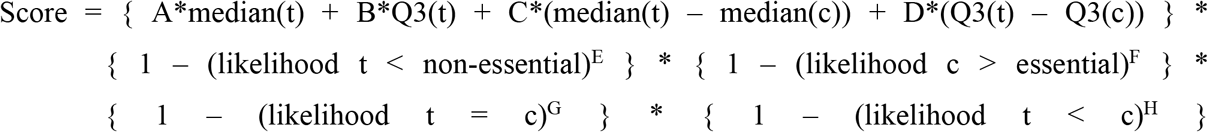

where A≅2, B≅0.017, C≅0.02, D=1, E≅8.8, F≅0.35, G≅7.1, and H≅0.22. Likelihoods are computed using Mann-Whitney U test where the inequality is tested by taking either the right or left tail and the equality is tested by taking both tails. For comparison with essential and non-essential genes, we use the gene sets described in ^40^(github.com/hart-lab/bagel). For essential genes, we use depletion values of all samples of all sgRNAs associated with an essential gene. For isogenic screens, we subtract 10,000 from all genes whose median(t) is less than zero. The top 3000 CCA scores are modeled using a beta distribution fitted using the fitdistrplus package ^41^ in R. Taking the top genes with p < 0.05, we stratify them into 4 Jenks classes using the classInt package in R (cran.r-project.org/web/packages/classInt/index.html). The values of the parameters, A through H except D, were determined by using a derivative-free optimiziation method, BiteOpt (github.com/avaneev/biteopt), to minimize:

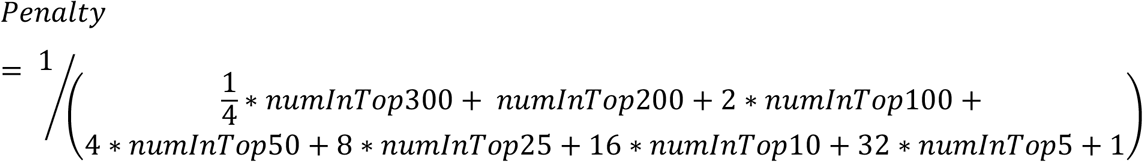

where numInTopN is the number of positive control synthetic lethal genes found in the top N genes as ranked by CCA’s scoring method over all training screens that have positive controls. D is always set to 1, as the other variables, A, B, and C, may be scaled. For the purpose of screens presented in this work, we considered a gene a hit if it is present in the top two Jenks classes.

### Public Cancer Dependency Data

Cell line panel estimates of gene dependency based on CRISPR screens were used in the analysis. CERES scores were downloaded from the 2020 Q1 release of the Broad Cancer Dependency Map (https://depmap.org/portal/download/). Copy Number Bias Corrected Fold Change Values were downloaded from the April 2019 release of the Sanger Project Score (https://score.depmap.sanger.ac.uk/downloads). The following cell lines were classified as BRCA1/2 biallelic mutants: COV362_OVARY, DOTC24510_CERVIX, HCC1395_BREAST, HCC1599_BREAST, ICC15_BILIARY_TRACT, JHOS2_OVARY, JHOS4_OVARY, MDAMB436_BREAST, SUM149PT_BREAST, CAPAN1_PANCREAS, SUM1315MO2_BREAST and UWB1289_OVARY. See Supplementary Table 4 for values.

### Antibodies

The antibodies listed below were used for immunoblotting (IB) or immunofluorescence (IF). Primary antibodies: mouse anti-CIP2A (clone 2G10-3B5; Santa Cruz sc80659, 1:500 IF, 1:1000 IB), rabbit anti-CIP2A (Cell Signalling Technologies #14805, 1:5000-12000 IB), rabbit anti-phospho-Histone H2A.X (Ser139) (Cell Signalling Technologies #2577, 1:500 IF), mouse anti-phospho-Histone H2A.X (Ser139) (clone JBW301; Millipore Sigma #05-636, 1:5000 IF), mouse anti-CHK1 (Santa Cruz sc8408, 1:500 IB), rabbit anti-phospho-CHK1 (Ser345) (Cell Signalling Technologies #2348, 1:1000 IB), rabbit anti-KAP1 (Bethyl A300-274A, 1:10000 IB), HRP-conjugated mouse anti-FLAG M2 (Sigma A8592, 1:1000-5000 IB), mouse anti-FLAG M2 (Sigma G1804, IB 1:1000), rat anti-FLAG (BioLegend #637301, 1:1000 IF), rabbit anti-TOPBP1 (Abcam ab2402, 1:2000 IF, 1:5000 IB or 1:1500 IB using yeast extracts), rabbit anti-TOPBP1 (ABE1463, Millipore, 1:300 IF), mouse anti-alpha-tubulin (Calbiochem CP06, 1:2000 IB), rat anti-HA (Roche 11867423001, 1:200 IB and IF or 1:500 IB using yeast extracts), mouse anti-CENPA (Abcam ab13939, 1:2000 IF), sheep anti-MDC1 (Serotec/Bio-Rad AHP799, 1:1000 IF), rabbit anti-MDC1 (Abcam ab11171, 1:1000 IF), rabbit anti-GAL4 DNA Binding Domain (Upstate 08283, 1:5000 IB), rat anti-Tubulin (YOL1/34) (Abcam ab6161, 1:2000 IB).

Secondary antibodies for immunoblots: IRDye 800CW goat anti-mouse IgG and IRDye 680RD goat anti-rabbit IgG (LiCOR 926-32210 and 926-68071, 1:5000 or 1:50000 using yeast extracts), HRP-conjugated sheep anti-mouse IgG (GE Healthcare NA931, 1:5000), HRP-conjugated goat anti-rabbit IgG (Cedarlane #111-035-144, 1:5000), and HRP-conjugated goat anti-rat IgG (Cedarlane 112-035-003, 1:5000 or 1:50000 using yeast extracts). Secondary antibodies for immunofluorescence: AlexaFluor 488-donkey anti-rat IgG (Thermo Fisher Scientific A21208, 1:2000), AlexaFluor 647-donkey anti-mouse IgG (Thermo Fisher Scientific A31571, 1:2000), AlexaFluor 488-goat anti-mouse IgG (Thermo Fisher Scientific A11029, 1:2000 or 1:1000 for high content microscopy), AlexaFluor 555-goat anti-mouse IgG (Thermo Fisher Scientific A21424, 1:2000), AlexaFluor 647-goat anti-mouse IgG (ThermoFisher Scientific A21236, 1:2000), AlexaFluor 647-goat anti-rabbit IgG (ThermoFisher Scientific A21244, 1:2000 IF), AlexaFluor 488-goat anti-rabbit IgG (Thermo Fisher Scientific A11034, 1:2000), AlexaFluor 555-donkey anti-sheep IgG (Thermo Fisher Scientific A21436, 1:2000), AlexaFluor 568-goat anti-rabbit IgG (Thermo Fisher Scientific A11011, 1:1000).

### Short interfering RNAs

The following siRNAs were used in this study: Dharmacon siGENOME Non-Targeting siRNA #3 D-001210-03-20, ON-TARGET Plus KIAA1524 (CIP2A) SMARTpool L-014135-01-0005, siGENOME MDC1 SMARTpool M-003506-04-0005, siGENOME TOPBP1 SMARTpool M-012358-01-0005).

### Fine chemicals

The following drugs were used in the course of the study: camptothecin (CPT, Sigma-Millipore), hydroxyurea (Sigma), nocodazole (Sigma), Shield-1 (Takara Bio USA, Inc), Aqua-Shield-1 (AS1; CheminPharma) and aphidicolin (Focus Biochemicals, 10-2058). Concentration and duration of treatment is indicated in the legends of the corresponding figures.

### High content imaging

To analyze γH2AX focus formation, cells were seeded in 96-well plates (~7,500 cells/well), cultured for 24 h, incubated in medium containing 20 mM EdU (5-ethynyl-2-deoxyuridine, Life Technologies) for the final 30 min and then washed with PBS and fixed with 4% paraformaldehyde (PFA) in PBS for 10 min at room temperature (RT). Cells were then processed for γH2AX staining. Prior to the click reaction, immunocomplexes were fixed again using 4% PFA/PBS for 5 min. Cells were rinsed with PBS and incubated with EdU staining buffer (150 mM Tris/HCl pH 8.8, 1mM CuSO4, 100 mM ascorbic acid and 10 mM AlexaFluor 647 Azide (Life Technologies)) for 30 min. After rinsing with PBS, images were acquired on an IN Cell Analyzer 6000 automated microscope (GE Life Sciences) with a 60X objective. Image analysis was performed using Columbus (PerkinElmer). Cell cycle profiling and analysis was evaluated based on EdU and DAPI staining.

### Immunofluorescence

Cells were grown and fixed on glass coverslips with 2-4% PFA, permeabilized with 0.3% Triton X-100 in PBS, and blocked with 5% BSA in PBS + 0.2% Tween-20. Cells were then stained for 2 h with primary antibodies in blocking buffer, washed three times with PBS + 0.2% Tween-20, incubated for 1 h with appropriate secondary antibodies plus 0.8 ug/ml DAPI, then washed twice with PBS + 0.2% Tween-20 and a final wash with PBS. Coverslips were mounted onto glass slides with ProLong Gold mounting reagent (Invitrogen). Images were acquired using a Zeiss LSM780 laser-scanning microscope (Oberkochen, Germany). Foci were manually counted.

For assessing the colocalization of MDC1, TOPBP1 and CIP2A, U2OS cells were reverse transfected with a final concentration of 10nM siRNA using Lipofectamine RNAiMAX (Invitrogen) on coverslips in 6-well plates. Nocodazole was added to the media at a final concentration of 100 ng/mL 16 h before collection. 48 h after transfection, cells were irradiated with 2 Gy of ionizing radiation using a Faxitron X-ray cabinet (Faxitron, Tucson, AZ) and allowed to recover for 1 h prior to fixation as described for immunofluorescence. Foci were counted manually and at least 50 mitotic cells per condition were imaged in each experiment.

For the experiments relating to the mitotic structures labelled by CIP2A and TOPBP1, U2OS wild-type and *MDC1^−/−^* cell lines were seeded on coverslips and either treated with 400 nM aphidicolin for 16 h (overnight) or left untreated. In order to perform immunofluorescence, cells were quickly washed once with cold PBS and then fixed with ice-cold methanol for 10 min on ice. Methanol was discarded and cells were washed two times with PBS before incubation with blocking buffer (10% FBS in PBS) for at least 1 h. Incubation with primary antibodies diluted in 5% FBS-PBS was performed overnight at 4°C in a humidity chamber. Coverslips were then washed 3 × 10 min with blocking buffer and incubated with AlexaFluor-conjugated secondary antibodies for 1 h at room temperature in the dark. After washing 3 × 10 min with PBS, coverslips were mounted on glass microscopy slides (Thermo Scientific, 630-1985, dimensions L76 × W26 mm) with VECTASHIELD mounting medium containing 0.5 μg/mL (DAPI) (Vector Laboratories, H-1200).

Confocal images were acquired using a Leica SP8 inverse confocal laser scanning microscope with a 63x, 1.4-NA Plan-Apochromat oil-immersion objective. The sequential scanning mode was applied, and the number of overexposed pixels was kept at a minimum. Images were recorded using optimal pixel size based on Nyquist criterion. At least 10 fields per condition with 10 to 15 z-sections were acquired, with 8-bit depth. Quantification of the foci was performed manually based on maximum intensity projections. Representative grayscale images were pseudocolored and adjusted for brightness and contrast in Adobe Photoshop CC 2020 by using adjustment layers.

### Immunoblotting

Cell pellets were boiled for 5-10 min in 2X SDS sample buffer (20% (v/v) glycerol, 2% (w/v) SDS, 0.01% (w/v) bromophenol blue, 167 mM Tris-Cl pH 6.8, 20 mM DTT) and separated by SDS-PAGE on gradient gels (Invitrogen). Proteins were transferred to nitrocellulose membranes (GE Healthcare), then blocked with 5% FBS or 5% milk in TBST and probed for 2 h with primary antibodies. Membranes were washed three times for 5 min with TBST, then probed with appropriate secondary antibodies for 1 h, and washed again with TBST, three times for 5 min. Secondary antibody detection was achieved using an Odyssey Scanner (LiCOR) or enhanced chemiluminescence (ECL, Thermo Fisher Scientific #34579).

### Cytogenetic analyses

To monitor chromosome aberrations, 0.5 × 10^6^ puromycin-selected RPE1-hTERT cells of the indicated genotypes were seeded in 10-cm dishes 3 d after transduction with virus particles expressing NLS-GFP-sgAAVS1 (control) or an NLS-GFP-sgRNA targeting a specific gene of interest. 4 d later 100 ng/mL KaryoMAX colcemid (Gibco/Thermo Fisher) was added for 2 h, and cells were harvested. To analyze sister chromatid exchange, 0.75 × 10^6^ RPE1-hTERT cells of the indicated genotypes were seeded in 10-cm dishes. 24 h after seeding, BrdU (final concentration 10 μM) was added to the media and cells were grown for 48 h; 100 ng/mL KaryoMAX colcemid (Gibco/Thermo Fisher) was added for the final 2 h. For cell harvesting, growth medium was stored in a conical tube. Cells were gently washed and treated twice for 5 min with 1 mL of trypsin. The growth medium and the 2 mL of trypsinization incubations were centrifuged (1000 rpm, 5 min, 4°C). Cells were then washed with PBS and resuspended in 75 mM KCl for 15 min at 37°C. Cells were centrifuged again, the supernatant was removed and cells were fixed by drop-wise addition of 1 mL fixative (ice-cold methanol:acetic acid, 3:1) while gently vortexing. An additional 9 mL of fixative was then added, and cells were incubated at 4°C for at least 16 h. Once fixed, metaphases were dropped on glass slides and air-dried overnight, protected from light.

To visualize chromosomal aberrations, slides were dehydrated in a 70%, 95% and 100% ethanol series (5 min each), air-dried and mounted in DAPI-containing ProLong Gold mounting medium (Molecular Probes/Thermo Fisher). To visualize sister chromatid exchanges (SCE) slides were rehydrated in PBS for 5 min and stained with 2 μg/mL Hoechst 33342 (Thermo Fisher) in 2xSSC (final concentration 300 mM NaCl, 30 mM sodium citrate, pH 7.0) for 15 min. Stained slides were placed in a plastic tray, covered with a thin layer of 2xSSC and irradiated with 254 nM UV light (~5400 J/m^2^). Slides were subsequently dehydrated in a 70%, 95% and 100% ethanol series (5 min each), air-dried and mounted in DAPI-containing ProLong Gold mounting medium (Molecular Probes/Thermo Fisher). Images were captured on a Zeiss LSM780 laser-scanning confocal microscope.

### LacR/LacO assays

For monitoring recruitment of endogenous CIP2A to FLAG-tagged TOPBP1 foci we used U2OS-FokI cells, which contain an integrated LacO array. These cells, which are known also as U2OS-DSB^42^, are referred to in the text as U2OS-lacO_256_ cells because we used them without any induction of FokI. 1.8 × 10^5^ cells were seeded in 6-well plates containing glass coverslips. 24 h after seeding, cells were transfected using 1 μg of pcDNA5-LacR-FLAG or pcDNA5-LacR-FLAG-TopBP1 (full length, fragments, or mutants) using Lipofectamine 2000. Cells were fixed with 4% PFA 48 h after transfection and stained for immunofluorescence. For monitoring recruitment of Flag-CIP2A, U2OS-FokI cells were transduced with pHIV-NAT constructs. After 0.1 mg/mL nourseothricin selection and cell expansion, 2 × 10^5^ cells were seeded in 6-well plates. The next day, cells were transfected using 1 μg of pcDNA5-LacR-TOPBP1. 24 h later, cells were seeded in a 96-well plate (~ 20,000 cells per well), cultured for 24 h and fixed with 2% PFA and stained for immunofluorescence. Images were acquired on an IN Cell Analyzer 6000 automated microscope (GE Life Sciences) with a 60X objective.

### Cell proliferation (IncuCyte) assays

MDA-MB-436, DLD1 wild-type and DLD1 *BRCA2*^−/−^ cells were infected with an empty virus containing the destabilization domain (DD) alone (pHIV-NAT-DD-FLAG) or virus containing an expression cassette for DD-tagged B6L (pHIV-NAT-DD-FLAG-TOPBP1-756-891). After nourseothricin selection (0.1 mg/mL for MDA-MB-436, 0.2 mg/mL for DLD1) and cell expansion, cells were seeded in 96-well plates (500-4,000 cells depending on cell line and genotype) and treated with 1 μM of Shield-1 or Aqua-Shield-1. The following day, plates were transferred into an IncuCyte Live-Cell Analysis Imager (Essen/Sartorius). Cell confluency was monitored every 4 h up to 10 d post-seeding.

### Micronuclei (MNi) assay

For TOPBP1 rescue experiments, DLD1 wild-type and *BRCA2*^−/−^ cells stably expressing 2xHA-TOPBP1 were generated by viral transduction and selection with blasticidin (7.5 μg/mL for parental cells, 10 μg/mL for *BRCA2*^−/−^ cells). 3 d after transduction with sgRNA viral particules (as described in the clonogenic survival assays), cells were seeded in a 96-well plate (1,500 for wild-type cells; 4,000 for *BRCA2*^−/−^ cells) and cultured for 4 additional days. For inducible B6L expression experiments, DLD1 and MDA-MB-436 cells were seeded in a 96-well plate (1,500 for DLD1 wild-type cells; 4,000 for DLD1 *BRCA2*^−/−^ cells; 14,000 for MDA-MB-436) and cultured for up to 4 days in the presence of Aqua-Shield-1. For detection of micronuclei, cells were fixed with 2% PFA, washed 3 times with PBS, permeabilized with 0.3% Triton X-100 in PBS for 5 min, washed 3 times with PBS, incubated for 1 h with PBS + DAPI (0.5 μg/mL). Alternatively, cells were stained for immunofluorescence (CENPA detection). After the last wash with PBS, images were acquired on an IN Cell Analyzer 6000 automated microscope (GE Life Sciences) with a 40X objective. Micronuclei were automatically detected and counted using the Columbus analysis tool (PerkinElmer).

### Yeast assays

Yeast two-hybrid assay was conducted using Matchmaker GAL4 two-hybrid system 3 (Clontech/Takara, USA). Bait and prey vectors were co-transformed into the yeast strain AH109 (Clontech/Takara, USA), using a standard high-efficiency transformation protocol, and plated onto media lacking tryptophan and leucine (SD-Trp-Leu) for 3 d to select for cells harboring the two plasmids. Single colonies were isolated and the interaction between bait and prey was assessed by a serial deletion assay based on the ability to grow on selective media lacking leucine, tryptophan, histidine and adenine (SD-Leu-Trp-His-Ade). Viability assays were performed using yeast cultures grown overnight at 30°C in SD-Trp-Leu to maintain plasmid selection. Ten-fold serial dilutions of cells were spotted on SD-Trp-Leu and SD-Leu-Trp-His-Ade containing 5 mM 3-amino-1,2,4-triazole (3-AT). Plates were imaged after 4 d of incubation at 30°C.

### Yeast protein extracts

For protein extracts, the cellular pellet of 20 mL of cell suspension (1×10^7^ cells/mL) was washed twice with 1 mL of 20% trichloroacetic acid (TCA) and suspended in 50 μL of 20% TCA. Cells were broken with acid-washed glass beads (Sigma G8772) by vortexing for 3 minutes at maximum speed. After addition of 100 μL of 5% TCA, precipitated proteins were transferred into a new 1.5 mL tube and centrifuged at 3000 rpm for 10 min at room temperature. The supernatant was removed, and the pellets of proteins suspended in 100 μL of 2X SDS sample buffer (20% (v/v) glycerol, 2% (w/v) SDS, 0.01% (w/v) bromophenol blue, 167 mM Tris-Cl pH 6.8, 20 mM DTT)). The pH was neutralized with 60 μl of 2M Tris base. The protein extract was boiled for 5 minutes at 95°C and centrifuged for 2 minutes at top speed at room temperature. The supernatant was collected, and the protein extract was subjected to SDS-PAGE analysis.

### Co-immunoprecipitation studies

Confluent 293T cells, either untreated or treated with 100 ng/mL nocodazole (Sigma) for 16 h, were used for each co-immunoprecipitation experiment. Cells were scraped directly into PBS, pelleted by centrifugation at 1000 × *g* for 5 minutes, and lysed by incubation in lysis buffer (50 mM Tris-HCl pH 8, 100 mM NaCl, 2 mM EDTA, 10 mM NaF, 0.5% NP-40, 10 mM MgCl_2_, 1x cOmplete EDTA-free Mini EDTA-free protease inhibitor tablet (Sigma), 1x Phosphatase inhibitor cocktail 3 (Sigma) and 5 U/mL benzonase (Sigma)) for 30 min on ice. Lysates were then cleared by centrifugation at 21,000 × *g* for 10 min. 1 μg of either mouse anti-CIP2A (2G10-3B5; Santa Cruz sc-80659) or normal mouse IgG (EMD Millipore 12-371) were added to the lysate and incubated with rotation at 4°C for 1 h. Subsequently, 20 μL of a slurry of protein G Dynabeads (Invitrogen) were added to the lysates and incubated for an additional 1 h at 4°C. Beads were collected using a magnetic rack and washed 4 × 5 min with 500 μL lysis buffer, then boiled in 25 μL 2X SDS sample buffer. The presence of co-immunoprecipitated proteins were detected by immunoblotting.

### Pharmacokinetic measurements

Whole blood was collected in over a period of 8 hr from conscious mice by tail snip and volumetrically transferred to tubes containing 0.1 M citrate (3:1 ratio blood/citrate) and frozen (−80°C). The determination of the total blood concentration was performed by protein precipitation extraction, followed by reversed-phase liquid chromatography and electrospray mass spectrometry (LC-MS/MS). Blood concentration versus time data was converted to plasma concentrations using an *in vitro* measurement of the blood to plasma ratio. The data were expressed as free plasma concentration using the fraction unbound which was assessed by equilibrium dialysis of AS-1 in mouse plasma over a period of 6 hours. PK profiles over a 24-hour period were estimated using Phoenix WinNonlin 8.3.1.

### Animals

Experiments were conducted in female NOD-SCID (Nonobese diabetic/Severe combined immunodeficiency) mice (5-7 weeks old, Charles River, St. Constant, Canada). Mice were group-housed on autoclaved corncob bedding in individual HEPA ventilated cages (Innocage® IVC, Innovive, San Diego, CA, USA) in a temperature-controlled environment (22±1.5 °C, 30-80 % relative humidity, 12-h light/dark). Mice were acclimatized in the animal facility for at least 5 d prior to use. Studies were conducted under a protocol that has been approved by Animal Care Committee. Animals were housed and experiments were performed at the Neomed site (Montreal, Canada), which has accreditation from CCAC (Canadian Council on Animal Care). Experiments were performed during the light phase of the cycle. Animals had irradiated food (Harlan Teklad, Montreal, Canada) and filtered water ad libitum. The number of animals used was the minimum necessary to achieve an 80% statistical power to detect a 40% change.

### Cancer cell implantation and measurement

DLD1 *BRCA2^−/−^* EV and B6L-expressing cells were harvested during exponential growth and re-suspended with high glucose RPMI1640 media (#30-2001, ATCC). Mice received a subcutaneous (SC) injection of 10×10^6^ DLD1 *BRCA2^−/−^* cells EV or B6L-expressing cells, in a volume of 0.1 μl, into the right flank. Tumor volume (TV) and body weight (BW) were measured 2-3 times per week. When tumors reached the target size of 150-200 mm^3^ mice were randomized into several groups (n=8) and treatment with AS-1 was initiated. Randomization was done to establish similar tumor volume mean and standard deviation in each group. AS-1 was administered Intraperitoneal (IP) twice daily (BID) in a volume of 5ml/kg in phosphate buffered saline (PBS). TV were measured using a digital caliper and calculated using the formula 0.52×L×W^2^. Response to treatment was evaluated for tumor growth inhibition (%TGI). TGI was defined as the formula: %TGI= ((TV_vehicle/last_ – TV_vehicle/day0_)-(TV_treated/last_ – TV_treated/day0_)) / (TV_vehicle/last_ – TV_vehicle/day0_)x100. BW is represented as change in BW using the formula: %BW change = (BW_last_ / BW_day0_)x100. Statistical significance relative to vehicle control was established by two-tailed unpaired Student’s t-test (Excel); *p<0.05; **p<0.01; *** p<0.001. All data are presented as the mean ± standard error of the mean.

---

**Fig. S1.**
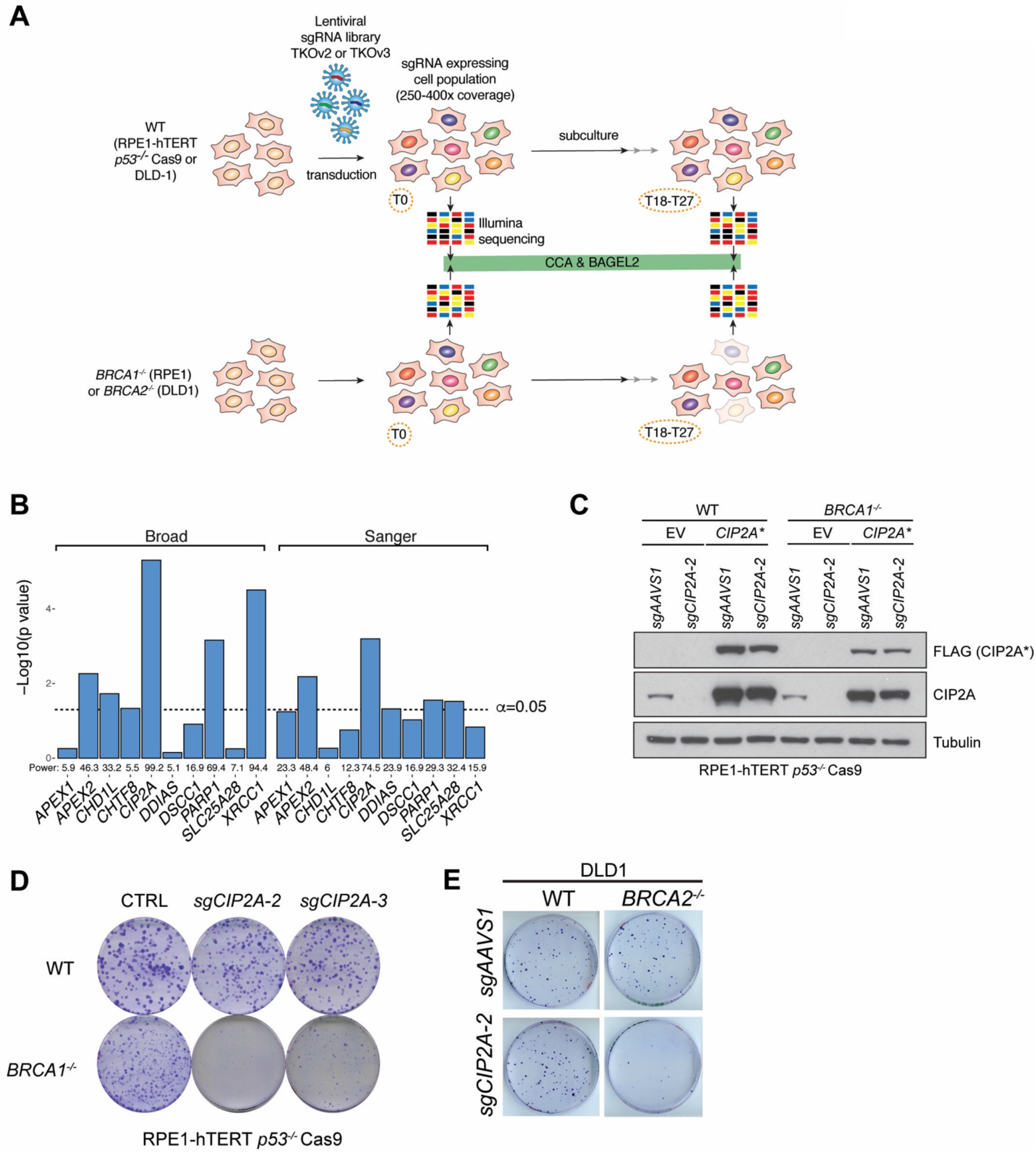
Supporting Data on the identification of *CIP2A* as synthetic lethal with BRCA1- and BRCA2-deficiency. (**A**) Schematic of the isogenic dropout CRISPR screens to identify synthetic-lethal interactions with BRCA1- and BRCA2-deficiency. (**B**) Statistical analyses for the data shown in Fig. 1BC. Shown are the results of a Mann-Whitney test comparing the values of the BRCA-proficient (BRCA^+^) and -deficient (BRCA^−^) gene depletion scores for the indicated genes. (**C**) Immunoblotting of whole-cell extracts of RPE1-hTERT *p53^−/−^* Cas9 cells, parental (WT) or *BRCA1^−/−^,* expressing the indicated sgRNAs and either a virus expressing an sgRNA-resistant *CIP2A* (*CIP2A\**) fused to a FLAG epitope-coding sequence or an empty virus (EV). Lysates were probed for FLAG (exogenous CIP2A), CIP2A and tubulin (loading control). (**D**, **E**) Representative images of the clonogenic survival assays shown in Fig. 1D (D) and the images for the DLD1 WT clonogenics relating to Fig. 1F (E).

**Fig. S2.**
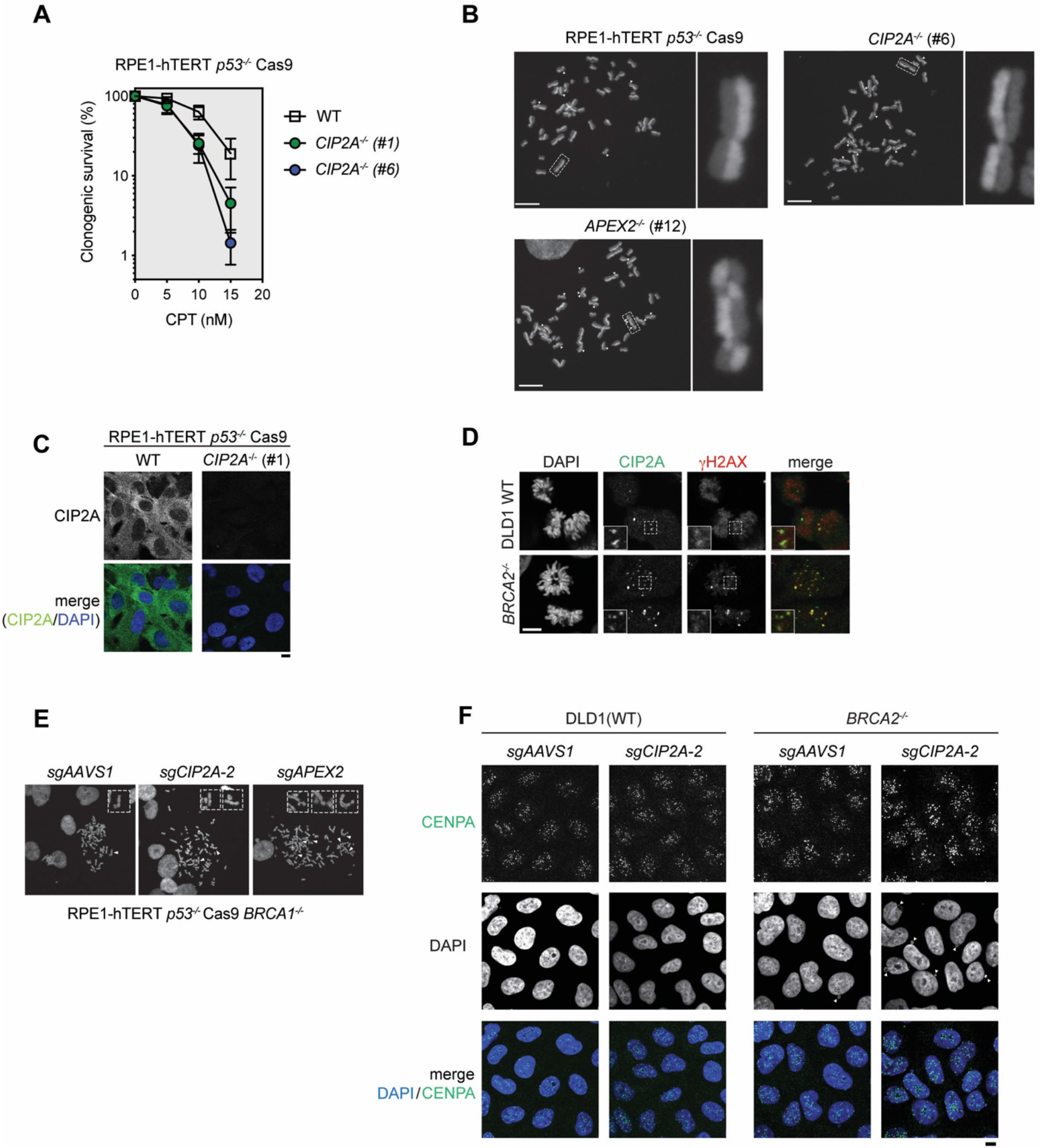
Loss of CIP2A does not cause DNA lesions requiring HR for their repair. (**A**) Clonogenic survival assay of RPE1-hTERT *p53^−/−^* Cas9 cells of the indicated genotype following treatment with camptothecin (CPT). Data points represent the mean ± S.D. (n=3). WT=wild type. (**B**) Representative micrographs of metaphase spreads for SCE analysis, relates to Fig. 2A. Arrowheads indicate an SCE event. Scale bar = 10 μm. **(C**) Immunofluorescence analysis of isogenic RPE1-hTERT *p53^−/−^* Cas9-derived WT and *CIP2A^−/−^* cells with a CIP2A antibody. Scale bar = 10 μm. (**D**) Representative micrographs of the experiment shown in Fig. 2G. Analysis of spontaneous CIP2A foci in DLD1 WT and *BRCA2^−/−^* mitotic cells. Scale bar = 10 μm. (**E**) Representative micrographs of the experiment presented in Fig. 2H showing scored radial chromosomes and chromosomes with chromatid breaks. Arrowheads indicate chromosome aberrations. (**F**) Representative micrographs of the experiment shown in Fig. 2I. White triangles show cells with micronuclei.

**Fig. S3.**
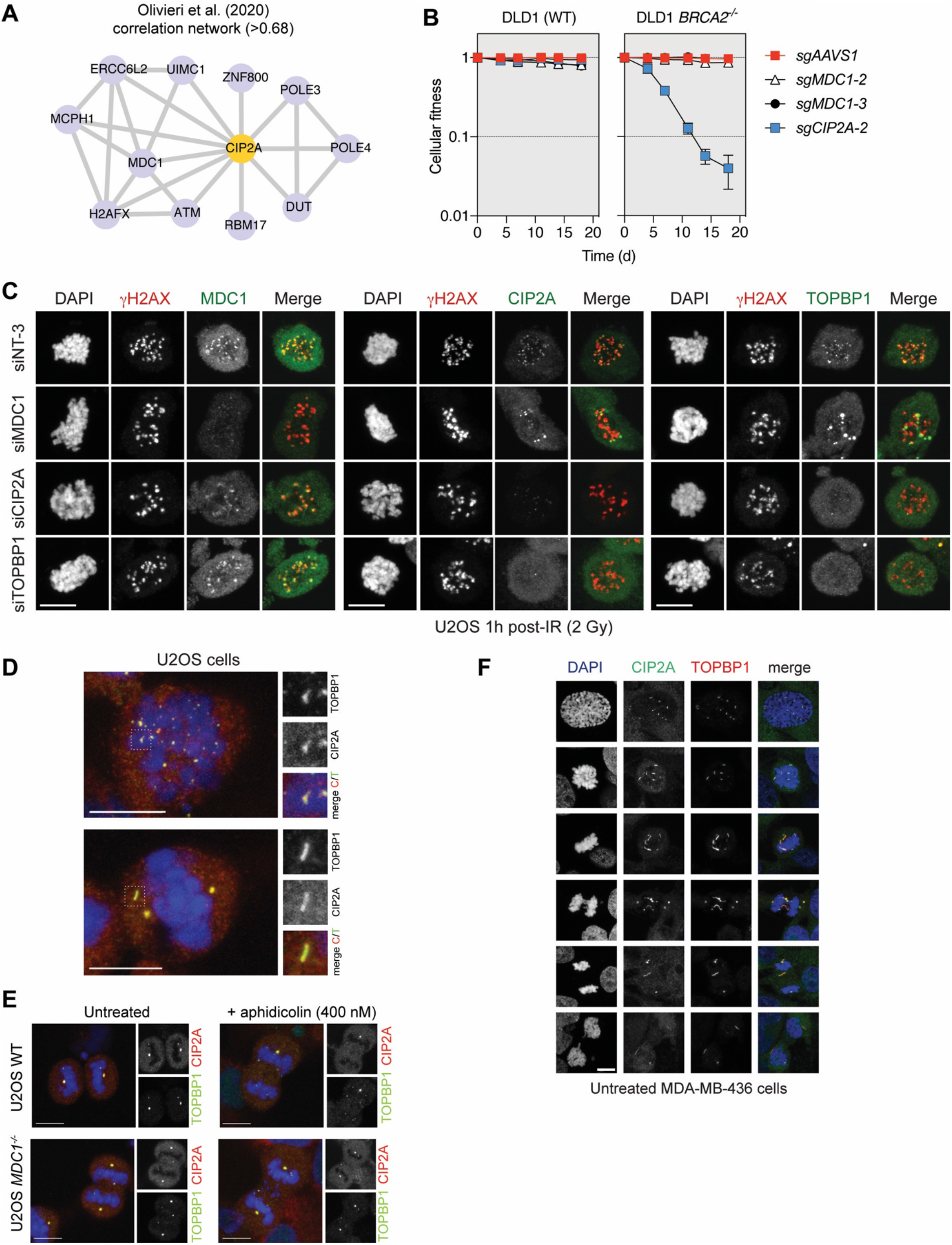
CIP2A acts in mitosis with TOPBP1. (**A**) Correlation network based on Pearson correlation of gene-level NormZ derived from the genotoxic dataset shown in ^21^. (**B**) Representative micrographs of the experiment quantitated in Fig. 3c. Nocodazole-treated U2OS cells previously transfected with either a non-targeting siRNA (siNT-3) or the indicated siRNAs were fixed 1 h post-X-irradiation (2 Gy) and processed for immunofluorescence with the indicated antibodies. Scale bar = 10 μm. (**C**) Competitive growth assays in DLD1 Cas9 (WT) or an isogenic *BRCA2^−/−^* counterpart transduced with virus expressing the indicated sgRNAs. Data are shown as mean ± S.D. (n=3 biologically independent experiments). (**D**) Additional micrographs of CIP2A/TOPBP1 structures observed in mitotic cells after treatment with low dose aphidicolin. Relates to Fig. 3E. Maximum intensity projections of confocal z-stacks. Scale bar = 10 μm. Shown here are curved (upper panels) and straight (lower panels) filaments. (**E**) Maximum intensity projections of confocal z-stacks of U2OS wild type and *MDC1*^−/−^ anaphase cells that were either treated with aphidicolin (400 nM) for 16 h or left untreated. Scale bars = 10 μm. Quantitation of this experiment is shown in Fig. 3F. (**F**) Representative micrographs of the experiment shown in Fig. 3G with additional MDA-MB-436 cells showing elongation of CIP2A-TOPBP1 filaments during mitosis. Maximum intensity projections of confocal z-stacks are shown. Scale bar = 10 μm.

**Fig. S4.**
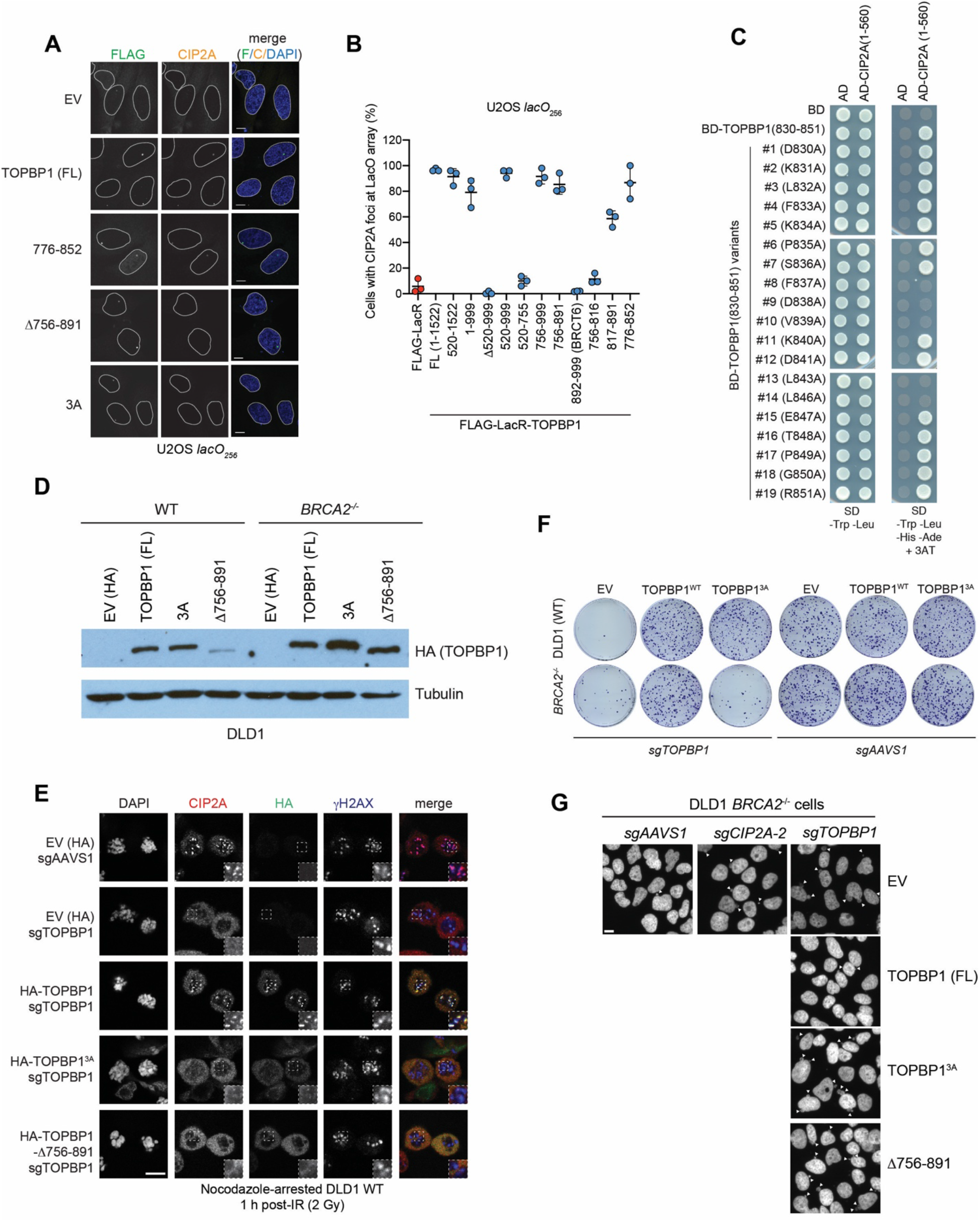
CIP2A interacts with TOPBP1 to promote BRCA-deficient cell viability. (**A**) Representative micrographs of the LacR/LacO assay assessing the interaction between endogenous CIP2A and TOPBP1 variants fused to Flag-LacR shown in Fig. 4F. Scale bars = 10 μm. (**B**) LacR/LacO assay assessing the interaction between endogenous CIP2A and TOPBP1 variants fused to Flag-LacR. Data points represent biological replicates, and the bars represent the mean ± S.D. (n=3). (**C**) Alanine scanning of TOPBP1 (830-851) residues by yeast two-hybrid with CIP2A (1-560). These studies identified 5 residues that abolish the TOPBP1-CIP2A interaction when mutated to alanine. AD=activation domain; BD=Gal4 DNA binding domain. Expression of proteins was verified by immunblotting but not shown. **(D**) Immunoblot of whole-cell extracts derived from DLD1 cells transduced with the indicated HA-tagged TOPBP1-expressing lentivirus or an empty virus that expresses an HA epitope (EV(HA)). The lysates were probed with an HA antibody or tubulin (loading control). FL=full-length. (**E**) Representative micrographs of DLD1 cells transduced with the indicated virus that were arrested in mitosis with a 16 h treatment with nocodazole, exposed to a 2 Gy IR dose and processed for immunofluorescence with the indicated antibodies 1 h later. Relates to the experiment quantitated in Fig. 4H. (**F**) Representative images of the crystal violet stains of the clonogenic survival experiment presented in Fig. 4I. (**G**) Representative micrographs of the experiment presented in Fig. 4J showing DAPI-stained cells to monitor micronucleation (labeled with arrowheads). Scale bar=10 μm.

**Fig. S5.**
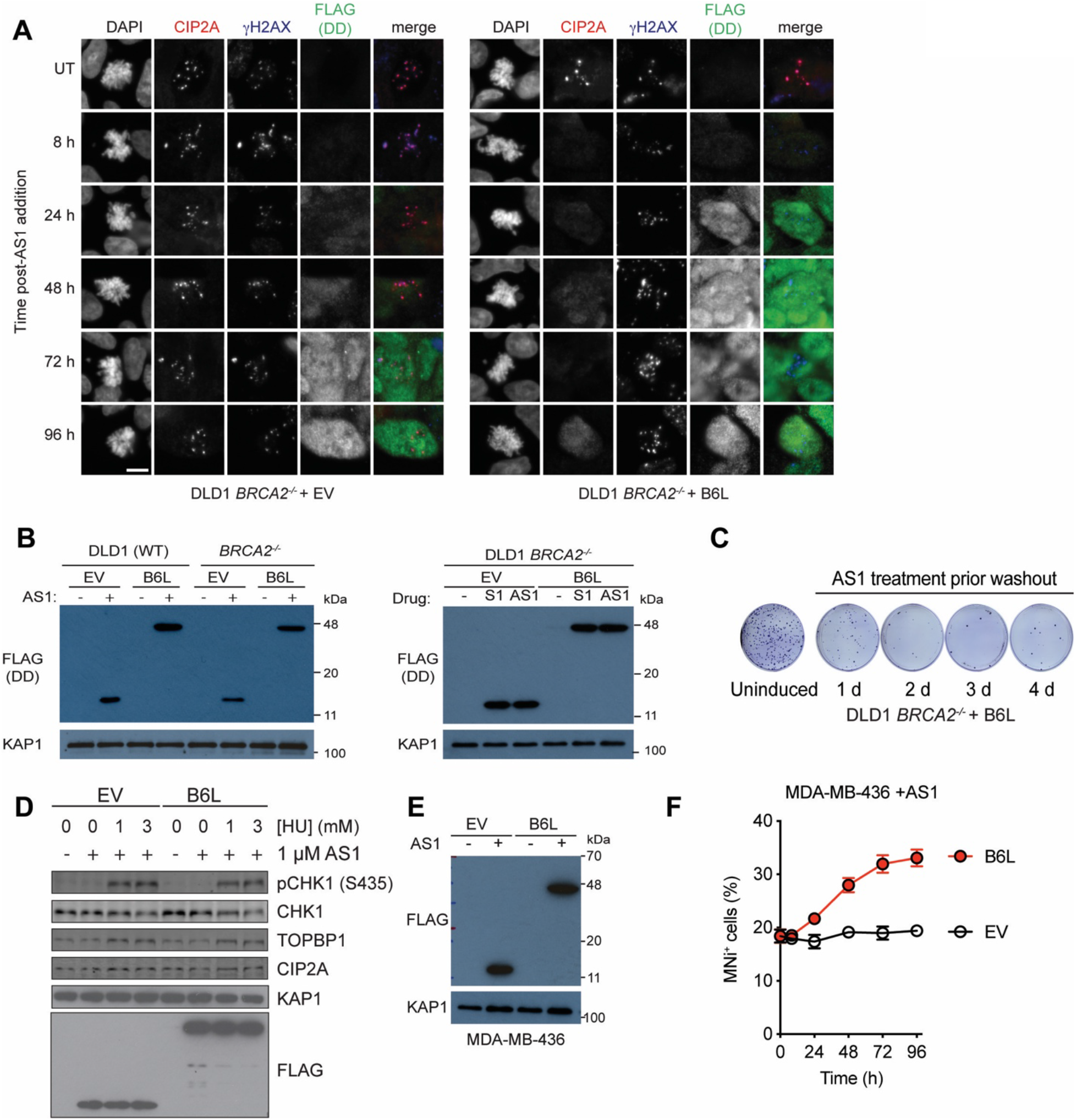
Disruption of the TOPBP1-CIP2A interaction is lethal in BRCA-deficient cells. (**A**) DLD1 *BRCA2^−/−^* cells transduced with either an empty virus containing only the destabilization domain (DD; EV, left) or a virus encoding B6L were treated with Shield-1 (1 μM) for the indicated periods or left untreated (UT). Shown are micrographs of mitotic cells stained for CIP2A, γH2AX or FLAG (labeling the DD). DNA was stained with DAPI. Scale bar = 10 μm. Quantitation of the experiment is shown in Fig. 5B. (**B**) Anti-FLAG immunoblots of whole-cell extracts derived from DLD1 parental (WT) or *BRCA2^−/−^* cells treated with either Shield-1 (S1) or Aqua-Shield-1 (AS1) for 72 h. These blots show similar induction of DD (in the empty virus; EV) or B6L upon addition of compound. Anti-KAP1 immunoblotting is used as a loading control. (**C**) Representative images of the clonogenic survival experiment presented in Fig. 5F. (**D**) Immunoblots assessing ATR signaling (CHK1 S345 phosphorylation) in DLD1 cells transduced with either an empty virus (EV) that expresses the unfused DD domain or a virus expressing B6L following induction with AS1. Cells were treated with hydroxyurea (HU) for the indicated times prior to harvesting. **e**, Anti-FLAG immunoblots of whole-cell extracts derived from MDA-MB-436 cells treated with Aqua-Shield-1 (AS1) for 72 h. Anti-KAP1 immunoblotting is used as loading control. (**F**) Quantitation of the of micronuclei (MNi)-positive cells in MDA-MB-436 transduced with an either empty virus (EV) or B6L-expressing virus following addition of AS1. Data is presented as mean ± S.D. (n=3).

**Fig. S6.**
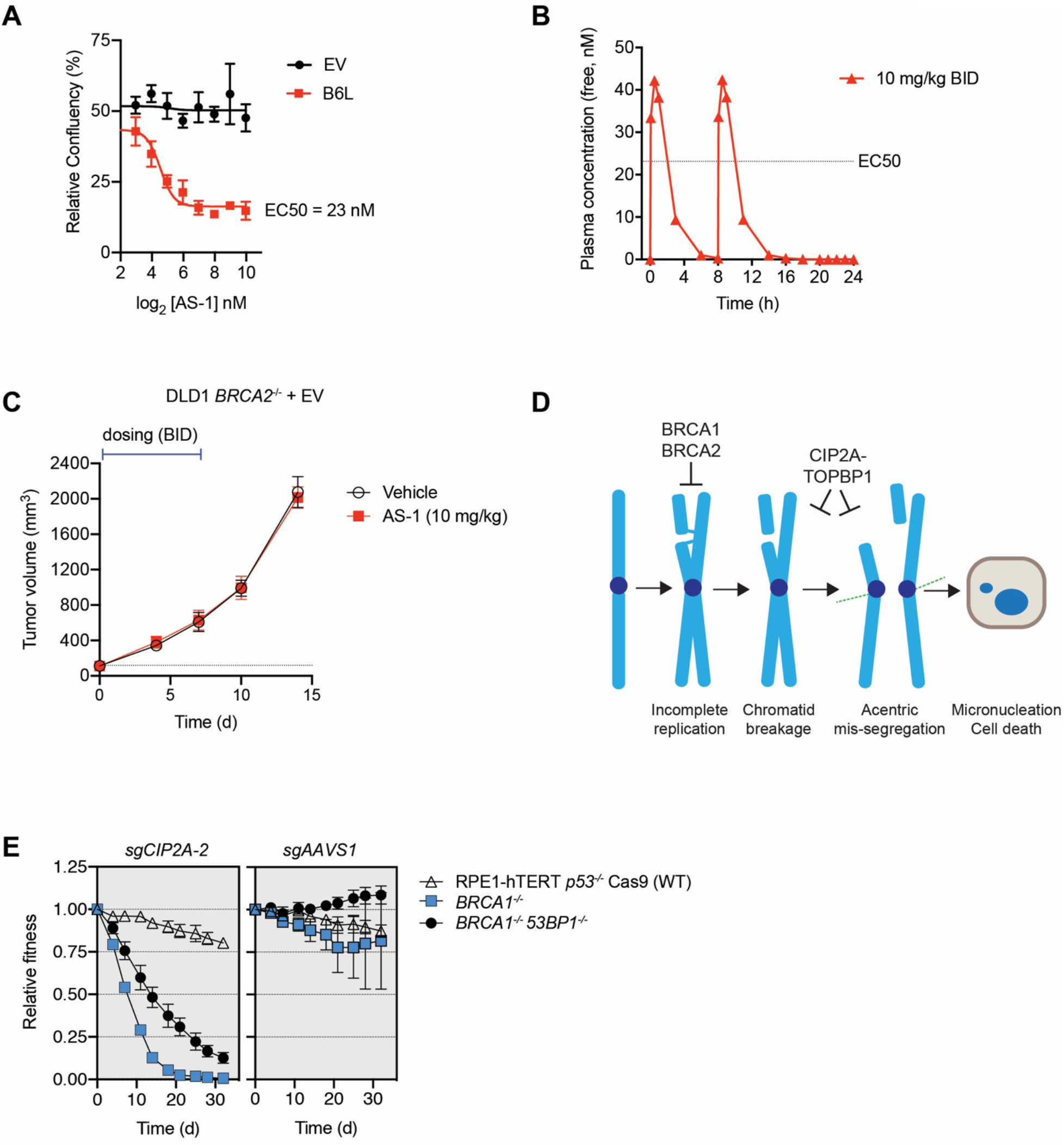
Control experiments to support the therapeutic hypothesis. **(A)** Determination of the EC50 growth inhibitory concentration of AS-1 in DLD1 *BRCA2^−/−^* cells transduced with lentivirus expressing either the DD domain alone (EV) or B6L. Growth was monitored in an Incucyte imager for 7 d. Data is shown as the mean ± SEM (n=3). (**B**) Pharmacokinetic analysis of AS-1 free plasma concentration over a 24 h period in mouse. Data is presented as the mean of values from 3 animals. (**C**) Growth of tumor xenografts derived from DLD *BRCA2*^−/−^ cells transduced with a DD-expressing lentivirus (EV) treated with AS-1 (20 mg/kg) intraperitoneally BID for 7d or with vehicle. Data is presented at the mean ± S.D. n=8. This acts as the control for Fig. 5J. (**D**) Model of the BRCA-CIP2A synthetic lethality. (**E**) Competitive growth assays in wild-type or RPE1-hTERT *p53^−/−^* Cas9 (WT) or isogenic *BRCA1^−/−^* or *BRCA1^−/−^ 53BP1^−/−^* counterparts transduced with virus expressing the indicated sgRNAs. Data are shown as mean ± S.E.M. (n=3 biologically independent experiments). Please note that the *53BP1*^−/−^ cell line was also subjected to transduction but is not shown for clarity.

**Table S1 (available as separate file).** Synthetic Lethality screen scores.

**Table S2 (available as separate file).** Raw values used to calculate plots in Fig. 1B,C

**Table S3.**
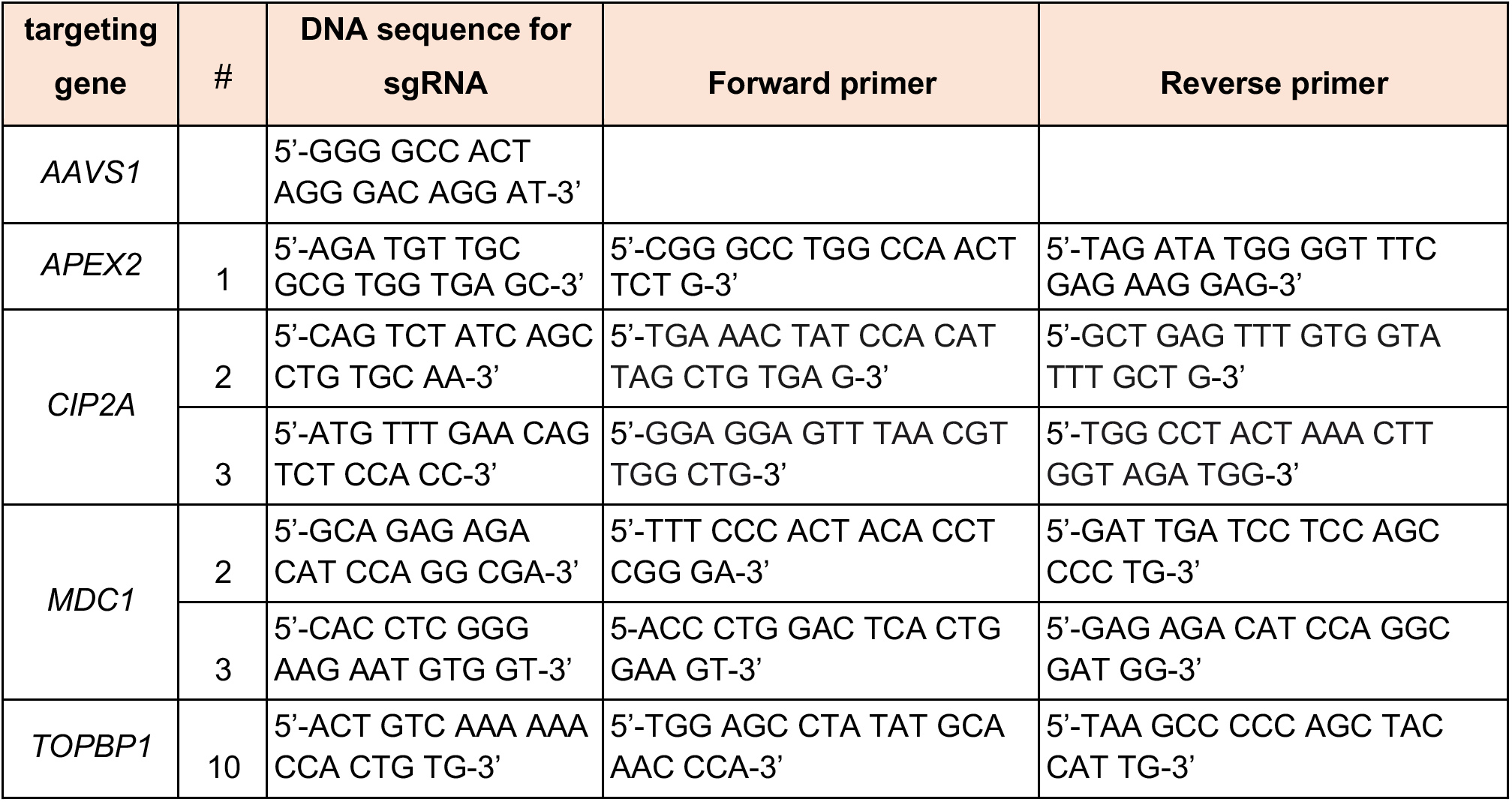
sgRNA used in the study along with primers used for sequencing the edited region in the genome (ICE analysis)

**Table S4.** Summary of the ICE editing analysis performed in the course of this study.

| Experiment | Cell line | Genotype | Other vector | sgRNA used | indel score (%) | KO score (%) |
| --- | --- | --- | --- | --- | --- | --- |
| Clonogenic survival assays |  |  |  |  |  |  |
| Related to Fig. 1D | RPE1 hTERT<br>Cas9 *p53*<sup>-/-</sup> | WT | NA | *CIP2A*-2 | 87 | 52 |
|  |  |  |  | *CIP2A*-3 | 64 | 51 |
|  |  | *BRCA1*<sup>-/-</sup> | NA | *CIP2A*-2 | 83 | 50 |
|  |  |  |  | *CIP2A*-3 | 69 | 55 |
| Related to Fig. 1F<br>+ 2I | DLD-1 | WT | NA | *CIP2A*-2 | 82 | 60 |
|  |  | *BRCA2*<sup>-/-</sup> | NA | *CIP2A*-2 | 60 | 46 |
| Rescue experiments |  |  |  |  |  |  |
| Related to Fig. 1E | RPE1 hTERT<br>Cas9 *p53*<sup>-/-</sup> | WT | EV | *CIP2A*-2 | 87 | 53 |
|  |  |  | Flag-*CIP2A* |  | 83 | 46 |
|  |  | *BRCA1*<sup>-/-</sup> | EV | *CIP2A*-2 | 87 | 52 |
|  |  |  | Flag-*CIP2A* |  | 80 | 50 |
| Related to Fig. 1G | DLD-1 | *BRCA2*<sup>-/-</sup> | EV | *CIP2A*-2 | 52 | 42 |
|  |  |  | Flag-*CIP2A* |  | 42 | 34 |
| Related to Fig. 4<br>H-J | DLD-1 | WT | EV | *TOPBP1*-10 | 90 | 90 |
|  |  |  | HA-*TOPBP1* |  | 96 | 96 |
|  |  |  | HA-*TOPBP1*<sup>3A</sup> |  | 98 | 98 |
|  |  |  | HA-*TOPBP1*-Δ756-891 |  | 95 | 95 |
|  |  | *BRCA2*<sup>-/-</sup> | EV |  | 81 | 81 |
|  |  |  | HA-*TOPBP1* |  | 87 | 87 |
|  |  |  | HA-*TOPBP1*<sup>3A</sup> |  | 92 | 91 |
|  |  |  | HA-*TOPBP1*-Δ756-891 |  | 75 | 75 |
| Two-color growth competition assay |  |  |  |  |  |  |
| Related to Fig. S3B | DLD-1 Cas9 | WT | NA | *CIP2A*-2 | 83 | 68 |
|  |  |  |  | *MDC1*-2 | 92 | 49 |
|  |  |  |  | *MDC1*-3 | 89 | 79 |
|  |  | *BRCA2*<sup>-/-</sup> | NA | *CIP2A*-2 | 60 | 46 |
|  |  |  |  | *MDC1*-2 | 74 | 53 |
|  |  |  |  | *MDC1*-3 | 63 | 60 |
| Related to Fig. S6E | RPE1 hTERT<br>Cas9 *p53*<sup>-/-</sup> | WT | NA | *CIP2A*-2 | 82 | 78 |
|  |  | *BRCA1*<sup>-/-</sup> |  |  | 86 | 75 |
|  |  | *BRCA1*<sup>-/-</sup><br>*53BP1*<sup>-/-</sup> |  |  | 91 | 56 |

## Dataset S1. (separate file)

*BRCA2*^−/−^ and *CIP2A*^−/−^ screen readcounts (TKOv3 library)

## Dataset S2. (separate file)

*BRCA1*^−/−^ screen readcounts (TKOv2 library)

